# Tuned inhibitory firing rate and connection weights as emergent network properties

**DOI:** 10.1101/2022.04.12.488114

**Authors:** Fereshteh Lagzi, Adrienne Fairhall

## Abstract

Excitatory cortical neurons show clear tuning to stimulus features, but the tuning properties of inhibitory neurons are ambiguous and have been the subject of a long debate. While inhibitory neurons have been considered to be largely untuned [1–4], recent studies show that some parvalbumin expressing (PV) neurons do show feature selectivity and participate in co-tuned subnetworks with pyramidal cells in which PV cells show high response similarity to the excitatory (E) neurons [5, 6]. Given shared input from layer 4 that drives feature tuning in excitatory subnetworks, we demonstrate that homeostatic regulation of postsynaptic firing rate governing the synaptic dynamics of the connections from PV to E cells, in combination with heterogeneity in the excitatory postsynaptic potentials (EPSP) that impinge on PV cells, results in the self-organization of PV subnetworks. We reconcile different experimental findings by showing that feature tuning of PV cells is an emerging network property that may be driven by synaptic heterogeneity, and can be inferred using population-level measures, while pairwise individual-level measures may fail to reveal inhibitory tuning. We show that such co-tuning can enhance network stability at the cost of response salience.

## Introduction

It is unclear how neurons in the cortex develop feature tuning. In mouse, excitatory neurons possess relatively strong feature tuning; however, there has been a long debate about the tuning of inhibitory neurons. Among different inhibitory subtypes in cortex which may play different and complementary roles in sensory processing [7, 8], Parvalbumin expressing subtypes (PV) play an important role in shaping the tuning responses of excitatory cells to different features of the sensory input [9–12]. Currently, there are two groups of seemingly conflicting studies about the tuning of PV cells to pyramidal excitatory (E) cells, both in terms of their firing rates and their patterns of connectivity to E cells in mouse V1.

One group of studies has reported that despite the tuning of the E cells to certain input features, such as orientation, individual PV cells are not tuned, or weakly tuned to those input features [1–4, 13], and therefore PV cells do not participate in co-tuned subnetworks with E cells. Given the dense connectivity profile of pyramidal excitatory to PV cells in layer 2/3 [2, 14], as well as the salt and pepper organization of feature maps in mice [15], PV cells might be expected to receive inputs from excitatory neurons tuned to many different features, remaining broadly tuned and without feature specific connections to excitatory cells.

A second set of studies, however, shows that despite the salt and pepper organization of feature maps in mice, PV cells can have a sharp tuning response [5]. Further, they can participate in subnetworks (motifs) of strongly coupled PV and E cells with relatively strong reciprocal connections [5, 6, 16]. Therefore, the claim is that in mice, PV tuning is related to the strength of the connections between E and PV cells, which correlates with the response similarity between PV and E cells [6]. In these reciprocal connections, there is a relatively constant EPSP to IPSP ratio [6] and this relation depends on the activity of the E and PV cells, and is mediated by layer 4 excitation [17, 18]. All of these studies, when put together, suggest that shared input, correlations between PV and E cells –and therefore feature specificity and the existence of co-tuned subnetworks of excitatory and inhibitory interactions– and the strength of their reciprocal connections are related to one another.

We reconcile these diverse experimental findings by showing in a model that the emergence of co-tuned PV neurons is a network property, and considering only pairwise correlations between PV and E cells may not identify their tuning. To achieve this, we consider two facts and one hypothesis about layer 2/3 neurons in mouse V1: (1) shared input from layer 4 drives feature tuning and stimulus selectivity in excitatory subnetworks [19, 20], (2) there is a large amount of heterogeneity in the excitatory postsynaptic potentials (EPSP) that impinge on PV cells [2, 14, 21], (3) we assume that the homeostatic regulation of postsynaptic firing rate which has been reported for GABAergic cells in the literature [22–25] governs the synaptic dynamics of the connections from PV to E cells. These considerations result in the emergence of co-tuned PV and pyramidal subnetworks, with a small portion of PV cells strongly tuned to Pyramidal cells. We show that PV tuning is a network property which may not be well identified with pairwise measures.

Our findings indicate that the higher the variance of the EPSPs of the connections from E to PV cells, the more tuned the PV cells become. Cotuned PV to E subnetworks provide stability to the network dynamics and expand the dynamic range of frequency responses of the E cells, but at the cost of reducing competition among excitatory assemblies, leading to reduced selectivity and reduced input amplification of the excitatory cells.

## Results

### Random connectivity from E to PV causes tuned PV to E connections

To study how PV cells develop their connections to E cells, we simulated three different networks with two excitatory assemblies (E_1_ and E_2_ in Fig. 1), and one population of PV cells. Neurons within each excitatory assembly had a stronger EPSP amplitude for their connections (*w J*, where *w* > 1), and received inputs from two sources: a common background source giving random and independent Poisson input to each neuron with a firing rate of (1 – *c*) *η,* and another shared correlated source with a firing rate of *cη,* which projected the same spike pattern to all neurons in each assembly, private to each assembly. In all networks, PV neurons were connected to each other and to the excitatory cells indistinguishably. Connections between PV cells had a constant IPSP amplitude, while connections from PV to E cells were plastic, following the symmetric STDP rule proposed in [22, 24].

**Fig. 1.**
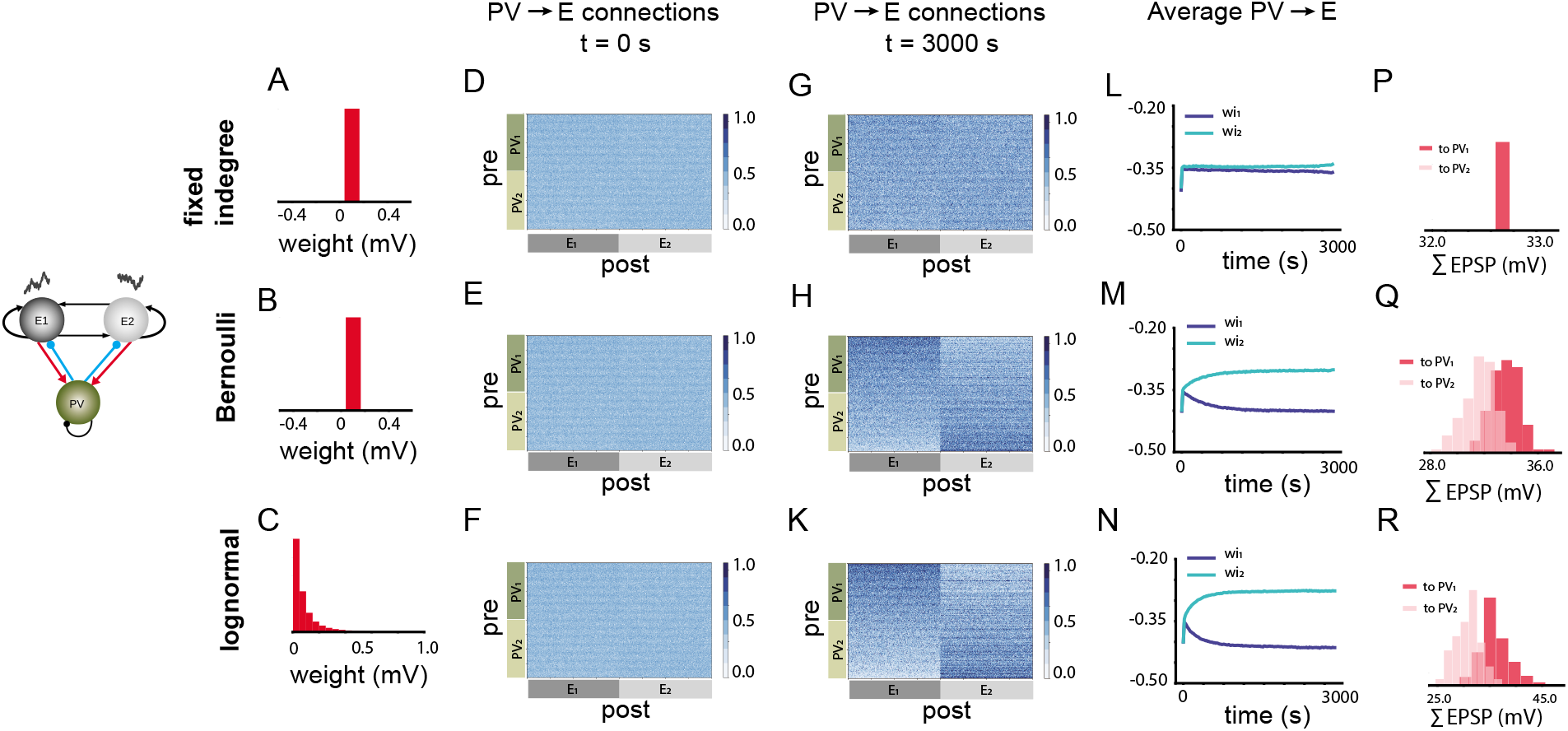
Effect of E to PV heterogeneity of connections on the emergence of tuned PV to E weights in a network with a single initial PV population. **A**: Fixed in-degree distribution of the weights with fixed EPSP amplitude. **B**: Existence of a connection between an E and a PV cell follows a Bernoulli distribution with a fixed EPSP amplitude. **C**: Connections between E and PV cells follow a Bernoulli distribution with EPSP amplitudes drawn from a log-normal distribution. **D-F**: Before initiating the plasticity rule, connectivity matrix for the connection weights from PV to E cells were originally random with a fixed IPSP amplitude. **G,H,K**: PV to E connectivity matrix at the end of the simulation, and PV index sorting for the networks with fixed in-degree, Bernoulli and log-normal weight distributions, respectively. PV_1_ and PV_2_ subnetworks are defined based on the block structure of the emerging connectivity matrix. **L, M, N**: Average IPSP amplitude for all connections from PV_1_ and PV_2_ neurons to E_1_ neurons. **P**: E_1_ to PV_1_ and PV_2_ summed EPSP distributions are the same due to the original fixed in-degree structure. **Q, R**: Distribution of total EPSP projections from individual E_1_ neurons to PV_1_ and PV_2_ neurons, after assigning memberships to PV cells.

In order to study the effect of heterogeneity in connections from E to PV cells, we chose different connectivity distributions, with identical mean values. The simplest network had a fixed in-degree distribution with identical EPSP amplitudes (Fig. 1A). The second network also had a fixed value of the EPSP amplitude; however, the probability of connections followed a Bernoulli distribution (Fig. 1B). Therefore, some PV cells receive more connections from E cells by chance. The third network also followed a Bernoulli distribution for the connections from E to PV cells; however, the EPSP amplitudes were drawn from a log-normal distribution (Fig. 1C).

For these different networks, we were interested in the evolution of the inhibitory weights over time. The IPSP amplitudes were initialized to be identical in all three networks (Fig. 1D, E, F). 2000 seconds after introducing the plasticity rule, we observed that the connectivity matrix denoting the absolute values of the IPSP amplitude from PV cells to E cells became structured for the networks with the Bernoulli (Fig. 1H) and log-normal distribution (Fig. 1K), but not for the network with fixed in-degree distribution. This can be seen by sorting the indices of the PV cells in the connectivity matrix according to their maximum total IPSP projection onto each excitatory assembly (Fig. 1G). The networks self-organized such that PV cells with relatively large values of ∑ | IPSP | for the outgoing connections to one excitatory assembly were more weakly connected, i.e. had smaller values of outgoing ∑ | IPSP |, to the other excitatory assembly. We then identified PV cells with stronger total connection weights to E_1_ (E_2_) as PV_1_ (PV_2_) cells. The average IPSP amplitude from PV_1_ to E_1_ grew as a function of simulation time for networks both with the Bernoulli (Fig. 1M) and the log-normal distribution (Fig. 1N). This growth was almost absent for the network with fixed in-degree distribution (Fig. 1L). These results indicate that for networks with heterogeneity in E to PV connections, a tendency for individual PV cells to develop stronger connections to some E cells was shaped during learning, and this preference was absent for the network with fixed in-degree.

To determine the reason for the emergence of this preference, we examined the differences in the distribution of the projections from each excitatory assembly onto the PV subnetworks for the different scenarios that we considered. More specifically, we calculated the distribution of the total (summed) EPSP from all E cells within E_1_ onto individual PV cells. For the networks with Bernoulli and log-normal distribution, we observed more distinct and skewed distributions of the total weights from E_1_ onto PV_1_ than from E_1_ onto PV_2_ (Fig. 1Q, R). However, these two distributions were identical for the fixed in-degree network (Fig. 1P). We confirmed that adding Hebbian E-to-PV and E-to-E plasticity results in the same pattern of PV weight evolution (Fig A1).

Taken together, we conclude that heterogeneity in the existence and the amplitude of the connections from E assemblies to PV cells can split the PV population into groups that by chance receive more projections from a certain excitatory assembly. The PV cells that received stronger total EPSP projections, connect to the corresponding excitatory assembly with stronger average IPSP values.

### Emergence of tuned PV to E weights and co-tuned PV subnetworks

To understand the circuit mechanisms underlying the emergence of tuned PV to E weights, we considered a simple network (Fig. 2A) with two excitatory populations, each received %30 of their inputs from a private shared input source. The network was also composed of two distinct populations of PV cells, each receiving stronger input from one of the excitatory assemblies. This was characterized by a factor *q* > 1 that scaled the stronger EPSP from one of the excitatory populations to a given PV population. The connections from the PV populations to the excitatory population that fed them with *q^J^* EPSP amplitude were determined by *w*_*i*1_, and the inhibitory connection weights to the other excitatory population were labeled as *w*_*i*2_. Initially, all IPSP amplitudes from the PV populations to the excitatory populations were identical.

**Fig. 2.**
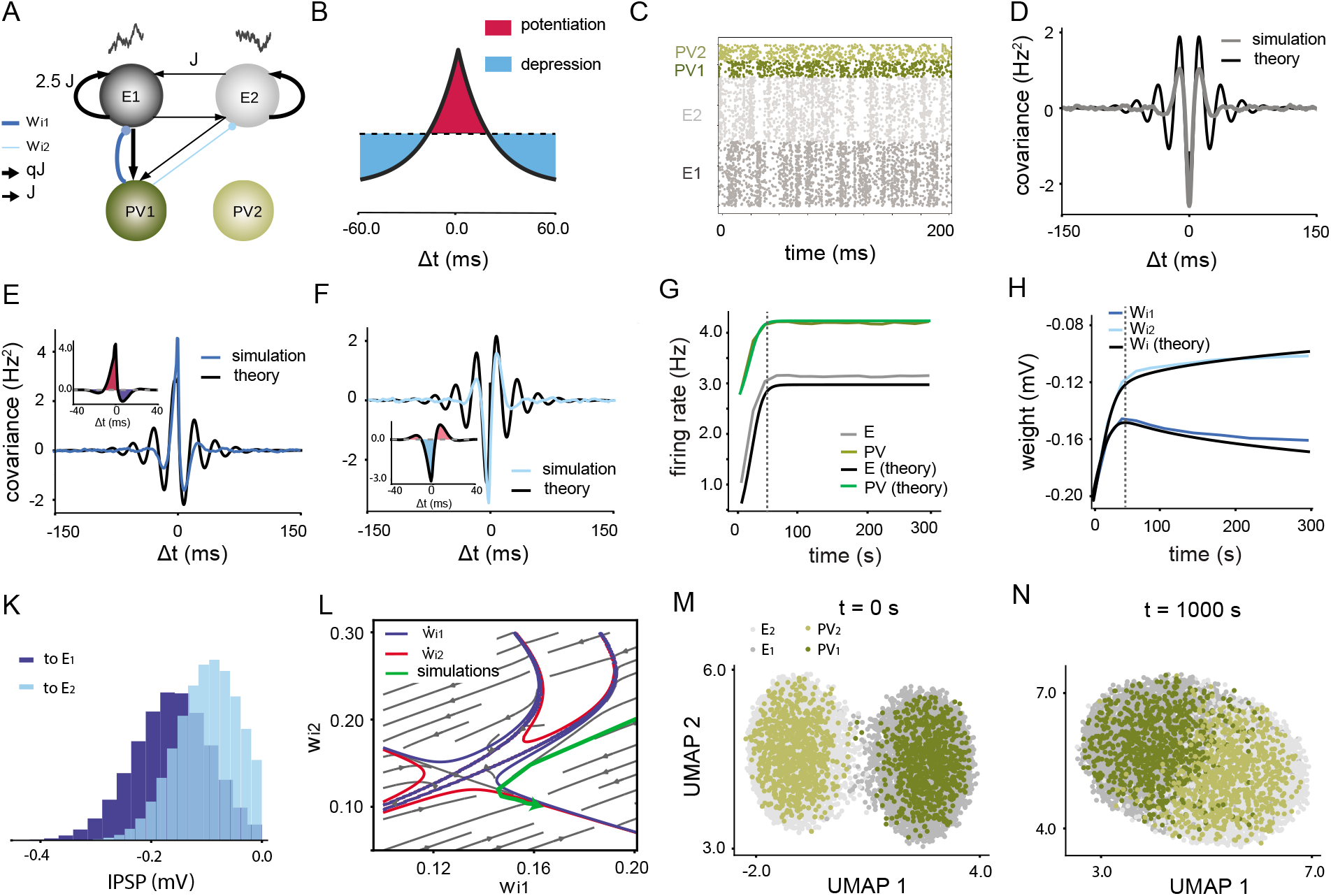
Theoretical understanding of the emergence of tuned PV subnetworks. **A**: Network with two excitatory and two inhibitory populations. **B**: Symmetric STDP curve which provides homeostasis for the excitatory firing rates. **C**: Raster plot of the network 1000 seconds after the onset of the inhibitory plasticity rule. E_1_ and PV_1_ are correlated (so are E_2_ and PV_2_). **D**: Covariance function between E_1_ and E_2_ average neuronal activity and the theoretical prediction for this function indicate that there is a strong competition between the two excitatory subnetworks. **E**: Covariance function between PV_1_ and E_1_ and the theoretical match to this function indicates a positive co-activation of the two populations. Inset: the learning signal has a net potentiation effect. **F**: Covariance function between PV_1_ and E_2_ and the theoretical match show a negative relationship and hence antagonism between the activities of E_2_ and PV_1_. Inset: the learning signal has a net depression effect. **G**: Firing rates of the excitatory and inhibitory populations reach the steady state value within a few 10 seconds, and our theory predicts the transient and steady state responses of the rates. **H**: Evolution of the IPSP weights from PV_1_ to E_1_ and E_2_ indicate a potentiation and depression, respectively, and our theory captures this phenomenon. **K**: Distribution of IPSPs from PV_1_ and PV_2_ onto cells in E_1_. **L**: Vector field of the coupled dynamics between average **w*_*i*1_* and **w*_*i*2_* derived from a linear theory. The trajectory of these weights obtained from large scale network simulations (green curve) follows the vector field and gets stuck at the intersection of the **w*_*i*1_* and *w*_*i*2_ nullclines, which indicates saturation of the weights. **M, N**: 2-dimensional UMAP plots for the raster plots of the network before and after plasticity.

The inhibitory plasticity rule applied (Fig. 2B) was similar to the model used in [24], consistent with reports from hippocampus [22], and also applied in recent findings on PV to E plasticity in OFC [25]. To justify the use of this plasticity rule in layer 2/3 in mouse V1, we refer to experimental findings in [26]. For a burst interval of 200 ms for a post synaptic excitatory neuron, it has been reported that the synaptic weight did not change when the pre-synaptic spike coincided with the first spike of the burst [26]. Considering the STDP function with a time constant of 20 ms (as observed in [22]), it would be rational to conclude that the STDP function had a potentiating role around small temporal differences (Δ*t* in Fig. 2B), and a depressing function for bigger Δ*t*. However, since the history of the burst for the postsynaptic neuron affects the result of the synaptic modification [27], the overall synaptic change on average would be around zero, as observed in [26].

The raster plot of the neuronal activities 1000 seconds after initiating the plasticity rule (Fig. 2C) shows that the two excitatory populations compete with each other. This competition can be well characterized by the average population cross-covariance function between the firing rates of the excitatory populations (Fig. 2D). We also observed strongly correlated activity between PV_1_ and E_1_ (similarly between PV_2_ and E_2_), and our mean-field model could capture the dynamics of this covariance very well (Fig. 2E, F). A positive covariance between PV_1_ and E_1_ as a result of strong EPSP for the connection from E_1_ to PV_1_ is apparent. Due to the symmetry in the connections, we expect a similar covariance function between PV_2_ and E_2_ (Fig. 2E). Also, since E_1_ and E_2_ are negatively correlated, the correlation between PV_1_ and E_2_ is mainly negative around zero-lag intervals (Fig. 2F).

In order to estimate the average inhibitory synaptic changes, we computed a “learning signal” (see Methods), for which a positive (negative) value indicates synaptic potentiation (depression). Because the average firing rates of the populations are identical for the presynaptic and postsynaptic ends of *w*_*i*1_ and *w*_*i*2_ (Fig. 2G), the only difference between the weight evolutions resides in the cross-covariance term. Since the learning signal is positive for *w*_*i*1_ (Fig. 2E, inset), this synapse is expected to grow as a function of the training time. However, the learning signal for the connection between PV_1_ and E_2_ is negative (Fig. 2F, inset), which results in an average depression of *w*_*i*2_ over time. The trajectory of the average inhibitory weights shows this separation (emergence of tuned weight) as a function of time (Fig. 2H). As a result of this emergence of tuned PV to E weights, the distribution of the IPSPs from PV_1_ to E_1_ is more skewed towards more negative amplitudes compared to the distribution of the projecting IPSPs to E_2_ (Fig. 2K).

The system of coupled rate and weight equations for the network in Fig. 2A can capture the dynamics of weight growth in the phase plane characterized solely by *w*_*i*1_ and *w*_*i*2_ (Fig. 2L). The majority of initial conditions in the region of interest shown in Fig. 2N will evolve toward a seeming line attractor formed by the intersection of the nullclines of 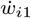 and 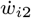 shown in blue and red at the bottom right corner of the state space. This line attractor is formed at higher values of *w*_*i*1_ relative to *w*_*i*2_, and hence explains why a reciprocally tuned inhibitory weight is formed in Fig. 2H. The trajectory which characterizes the evolution of the average inhibitory weights in our large scale simulations is shown in the phase plane (Fig. 2L) and its dynamics is well predicted by the flow of the system.

If shared correlated input is viewed as an external feature that correlates a group of neurons, then the development of reciprocal inhibition comes at the cost of diminished feature specificity in excitatory assemblies. The reason is that reciprocal inhibition decorrelates excitatory neurons and can potentially prevent input amplification. This can be shown by applying a 2-dimensional UMAP [28] to the neuronal activities, with cosine similarity distance measure (Fig. 2M, N). Before evolving through plasticity, the excitatory assemblies form well separated clusters, and due to strong correlations between E and PV cells, PV populations also cluster together with their corresponding E assemblies. However, after the operation of the inhibitory plasticity, as a result of developed reciprocal inhibitory weights, the excitatory clusters are not as well separated and become more intermixed, but the PV cells are still clustered with their own E assemblies. This indicates that the spontaneous activities of the E cells are less distinguishable and hence less susceptible to amplify the external input.

### Importance of internal correlations

In the emergence of tuned PV to E weights, two correlation terms play roles. First, shared *external* input provided to the E cells in each assembly, with a firing rate of *c η*; increasing the value of c increases this correlation term, and results in more separation between *w*_*i*1_ and *w*_*i*2_ (Fig. 3A). The second correlation term is generated *internally* between the excitatory and inhibitory neurons, due to direct wiring with different values of EPSP amplitudes. Even without any shared correlated input (with *c* = 0), a weak tuned PV to E weight structure emerged (Fig. 3A, light blue curves) resulting from internal network dynamics. For non-zero values of *c* this term plays a bigger role. To demonstrate the importance of this contribution in our theory, for *c* = 0.3, we compare with results obtained by ignoring all the contributing terms which characterized the internal correlations between PV and E cells. These terms had no impact on the steady state population firing rates (Fig. 3B), and simulation and theory converged to similar results. However, a dramatic effect was seen in the dynamics of the weight evolution (Fig. 3C): both *w*_*i*1_ and *w*_*i*2_ evolved similarly and as a result converged to identical values around the average value of the steady state weights for the simulation in Fig. 2H. This clearly does not match with the simulation results in Fig. 2H. We thus conclude that the internal correlations between the inhibitory and excitatory cells have a major contribution in shaping tuned PV weights. Ignoring those correlation terms changed the dynamics of inhibitory weight evolution inconsistent with the network dynamics.

**Fig. 3.**
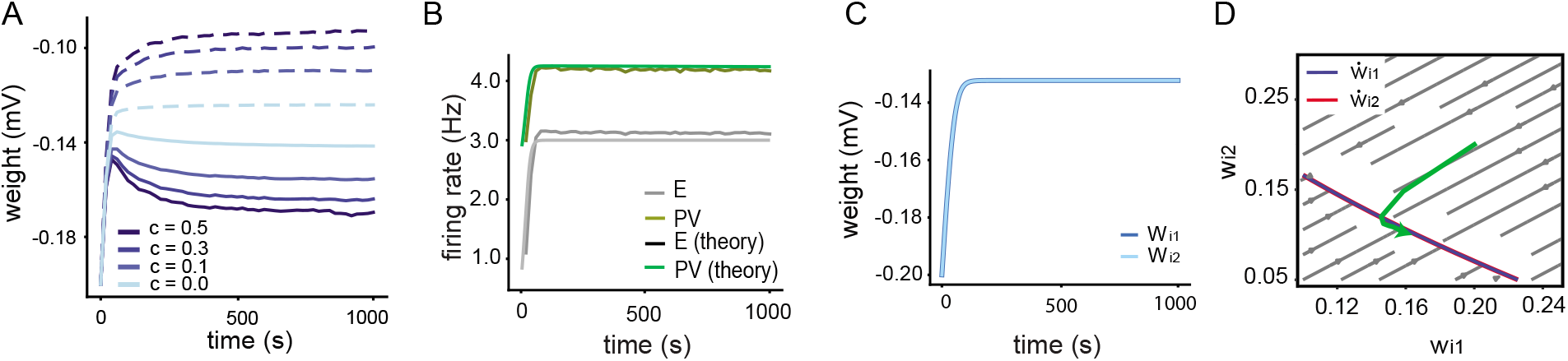
Importance of correlations and consequences of ignoring them. **A**: Increasing the percentage of input correlations *c* compared to the background input has a dramatic positive effect on the separation of the IPSP amplitudes to different excitatory assemblies. **B**: Ignoring the correlation term in the theoretical derivations does not affect the rate dynamics and our theory can capture the dynamics of the excitatory and inhibitory rates. **C**: The evolution of the IPSP amplitudes from PV_1_ to E_1_ and E_2_ would be exactly identical if the correlation term is removed from the equations. **D**: Vector field of the system describing *w*_*i*1_ and *w*_*i*2_ coupled dynamics. The nullclines of 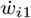 and 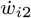 are exactly equal and their intersection forms a line attractor. Trajectory of the weight evolution from network simulations (green curve) does not match the flow of the vector filed.

Ignoring the correlation terms between PV and E cells in our theory resulted in a vector field with overlaid nullclines which formed a line attractor (Fig. 3D). Depending on the initial conditions of the average inhibitory weights from PV to E population, the steady state values of the weights may land on different points of the line attractor. For example, the weight trajectory from Fig. 3C, which shares the same initial condition with the network, moves on a straight line and converges to the closest point on the line attractor (blue trajectory in Fig. 3D). This result is not consistent with the trajectory from the simulation (green trajectory).

We conclude that even under spontaneous activity in which there are no modulations in the external firing rate, the internal dynamics of the network, particularly the correlations between the PV and E cells, determine the steady state values of the inhibitory weights which result in tuned PV to E weights. Ignoring the correlation term in our mathematical analysis results in identical inhibitory weights, without any emergence of tuning between PV and E.

### Monotonic relationship between EPSP and IPSPs and its role in response tuning

Previously, we showed that heterogeneity in EPSP distribution for the connections from E to PV cells can result in different degrees of tuned PV to E cells in different network scenarios (Fig. 1). We now explore patterns of the reciprocal EPSPs and IPSPs when the network is composed of multiple subnetworks (more than 2 excitatory assemblies) and investigate how these patterns affect tuning of PV and E responses.

We considered a network with four excitatory assemblies (E_1_ through E_4_), and assigned four distinct PV populations (PV_1_ through PV_4_) (Fig. 4A). We considered a network with a large amount of heterogeneity for the E to PV connections (wide network), and another network with relatively small heterogeneity for those connections (narrow network). The EPSP values from E_*x*_ to PV_*x*_, *x* = 1,…,4 was the strongest (characterized by *q J*, with *q* = 2.5 for the wide network and q = 1.85 for the narrow network), and decreased from E_*x*_ to PV_*x*+*n*_ in a cyclic manner. We also consider the delta network, which is the extreme case of the narrow network with no variability between the EPSP to PV weights (all q values are identical and equal to 1.75).

**Fig. 4.**
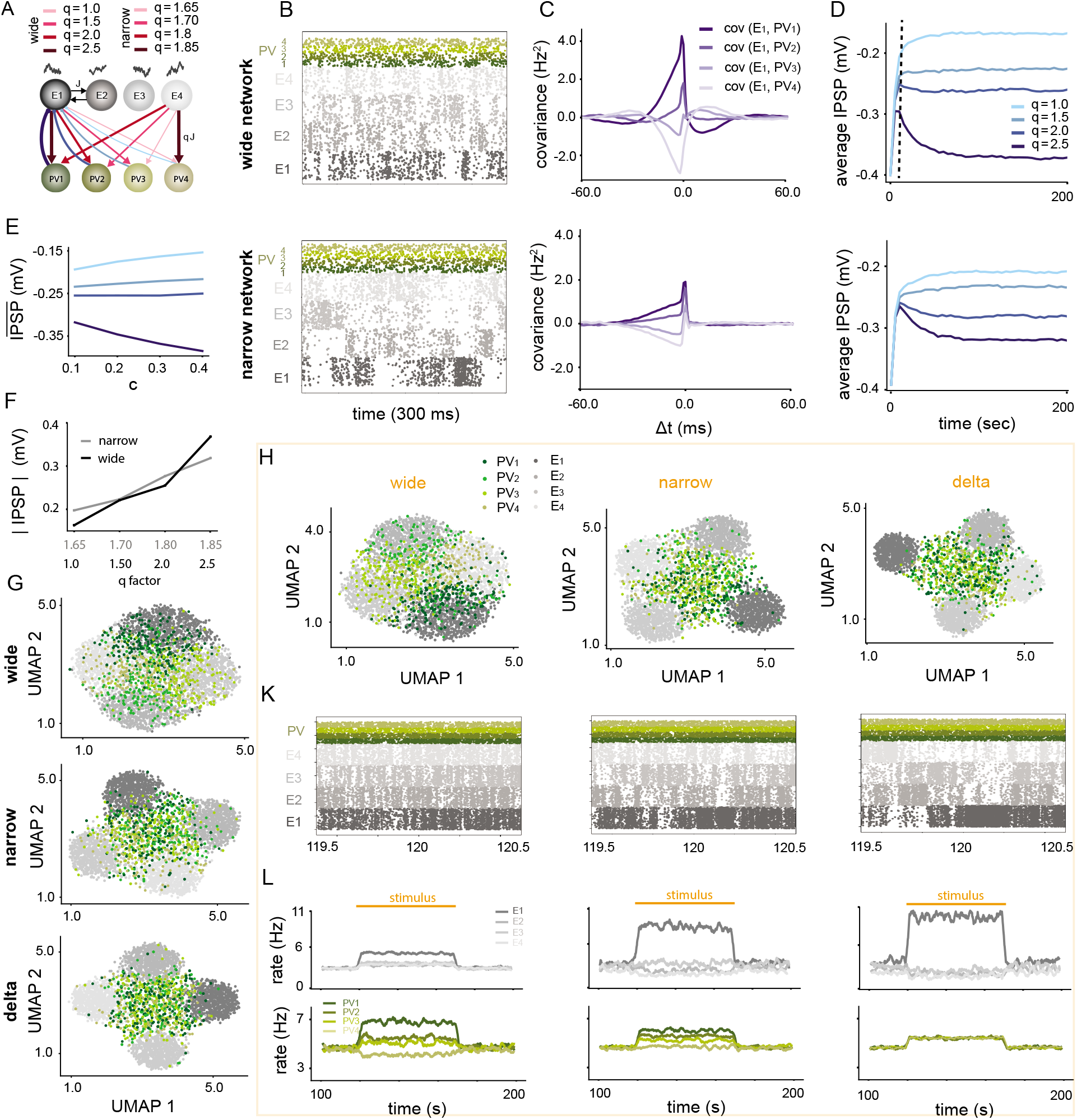
Firing rate homeostasis provided by inhibitory plasticity results in a monotonic relationship between reciprocal EPSP and IPSP amplitudes and reduces feature selectivity of the network. **A**: Network with 4 similar excitatory assemblies and a profile of connectivity to PV subnetworks with varying EPSP amplitudes modified by (q = 1.0, 1.5, 2.0, 2.5) for the wide network and by (q = 1.65, 1.70, 1.80, 1.85) for the narrow network. All reciprocal IPSP amplitudes are identical at the beginning of the simulation. **B**: Raster plot for the wide network (top) and narrow network (bottom) after 1000 seconds of training indicates high correlations between excitatory and their highly tuned PV assembly. **C**: Covariance function between average PV_1_ and average E_1_ neuronal firing rates for the wide network (top) and narrow network (bottom).**D**: Evolution of IPSP amplitudes from PV_1_ through PV4 onto E_1_ assembly. Smaller values of *q* in the corresponding EPSP amplitude from E_1_ results in weaker reciprocal projection from the PV subnetwork back onto E_1_. **E**: Steady state amplitude of the IPSP corresponding to different reciprocal q value for the EPSP of the projecting connection as a function of increasing the value of input correlation c. The distance between the IPSP amplitudes increases with increasing c. **F**: Monotonic relationship between IPSP and EPSP amplitudes as a function of increasing q for the wide (black) and narrow (gray) network. **G**: From top to bottom: 2-dimensional UMAP plots for the neuronal activities for the wide, narrow and delta networks show more separable activities for the E cells, but less tuned PV to E neuronal activities. **H**: From left to right: 2-dimensional UMAP plots for the wide, narrow and delta network when E_1_ receives specific sensory input. **K**: Raster plots of the networks before and after the onset of sensory input to E_1_ at t = 120s. **L**: Average neuronal firing rates for the excitatory (top) and PV (bottom) populations in 100 seconds interval for which sensory input evokes E_1_ activity from t = 120 s to t = 170 s.

After 1000 seconds of simulation time, shared input together with the plasticity rule resulted in a raster plot with correlated *PV_x_* with *E_x_* activity, and correlated activities of the E cells within each assembly (Fig. 4B). Due to the symmetry of the pattern of connections from the excitatory assemblies to the PV populations, the average firing rates of all excitatory cells were identical. For a similar reason, the average of the firing rate for all PV populations also reached an identical steady state value (Fig. A2B).

Strong projections from excitatory assemblies to individual PV populations resulted in a strong positive covariance between the two populations (Fig. 4C). Due to stronger EPSP projections in the wide network, the amplitude of those correlation functions were bigger (Fig. 4C top compared to bottom). However, the weakest projection from E_1_ to PV_4_ caused a negative covariance between the two populations. The average inhibitory weights from PV_1_ through PV_4_ to E_1_ reflect a pattern similar to the strength of the projecting EPSP from E_1_ to the four PV populations (Fig. 4D). In other words, the value of the IPSP from PV_1_ to E_1_ is the strongest in absolute value since PV_1_ received the strongest EPSP (*q* = 2.5) from E_1_, which then resulted in a strong covariance between the two populations, and hence increased the learning signal in Fig. A2C. The monotonicity in the strength of average EPSP and IPSP between populations is clearly visible in Fig. 4D. The difference between the average IPSP amplitudes is smaller for the narrow network (Fig. 4D, bottom) due to weaker covariances between PV and E assemblies (Fig. 4C). For increasing values of the external correlated input (*c*), the difference between the IPSP amplitudes at the steady state increases (Fig. 4E) because increasing correlations in the external input strengthens the covariance between PV and E assemblies. This indicates that the PV connections to the excitatory assembly which feeds them strongly potentiates, and the connections to those assemblies which feed them weakly depresses. This results in a monotonic relationship between the reciprocal EPSP and IPSP values both in wide and narrow networks, however, the variance of the IPSP projections is bigger for the wide network (Fig. 4F).

### Tuned PV to E subnetworks reduce feature tuning of the E cells

We compared the separability (and hence selectivity to input features due to correlated input) of the neuronal activity patterns for the wide, narrow and delta network in the low dimensional UMAP space, both for the spontaneous (Fig. 4G) and evoked (E_1_ received sensory input, Fig. 4H) states. For all networks in both spontaneous and evoked conditions, excitatory neurons within each assembly had the most similar (and hence correlated) patterns of activity, while excitatory neurons in different assemblies had distinct activity patterns and formed distinct clusters. Due to the strong EPSP projections and high covariances between the E and PV populations in the wide network, PV neuronal responses also formed distinct clusters around their corresponding E assemblies from which they received strong excitatory inputs. This indicates the emergence of feature tuning for the PV cells in the wide network due to internal correlations, although the PV cells had not received any external correlated input. For the narrow network, due to smaller variance of projections from E to PV cells, PV neuronal activities were much less tuned to the excitatory assemblies, and did not form distinct clusters. The extreme case of the narrow network, the delta network, resulted in a maximally separate pattern of E activities for the different assemblies, however, the PV neuronal dynamics were not tuned to the excitatory responses, and also no distinct clusters of PV activities were formed (Fig. 4G, H). The reason is that in this case, the covariance functions between the PV population activities and the excitatory assembly firing rates are all identical (Fig. A2G).

In the spontaneous activity condition we have modeled so far, distances between the E clusters for the wide network were not as large as those for the narrow and delta networks (Fig. 4G). This is related to the weak competition between the excitatory assemblies (Fig. A2D) as a result of strong reciprocal inhibition provided by PV populations which decorrelates the excitatory activities and tends to homogenize the activities of the E cells.

We next presented an external specific input to E_1_ cells and a smaller nonspecific and common input to all excitatory clusters. As a result of this input, for all networks, the E_1_ cluster was pushed away from the rest of the E clusters (Fig. 4H,K,L). However, the distance between E_1_ and the rest of the clusters was minimum for the wide network compared to the narrow and delta networks. Therefore, feature selectivity for the wide network is relatively weak. Formation of more separate clusters for the narrow and delta networks is related to stronger competition between the assemblies (Fig. A2E, F compared to D). Less variable projections from E to PV create weaker reciprocal inhibitory connections back onto the E assemblies, and hence cause less decorrelation between the E cells, which results in larger distance between the E clusters in the UMAP plots.

These inferences that we have drawn based on distances in the lowdimensional UMAP space are supported by the firing rate dynamics of the network. The raster plots before and after the onset of stimulus at t=125 s are more distinct for the delta case compared to other networks (Fig. 4K). Also, the average firing rates of the excitatory cells show a big input amplification for the delta network compared to the baseline activity. This amplification is minimal for the wide network. Since in the wide network case PV cells are more strongly tuned to the E cells, PV_1_ activity is also amplified as a result of E_1_ increased firing rate (Fig. 4L, bottom row), and represents feature selectivity due to its tuned response to E_1_ activity. This amplification is reduced for the narrow network, and finally for the delta network there is no separation between PV firing rates. This results from the lack of tuning between PV and E cells in this case.

To summarize the results of this section, first we highlight how diversity and heterogeneity in the excitatory connections onto PV cells result in stronger reciprocal inhibitory projections from the PV populations. Second, we show that the larger the variance of the excitatory connections onto the PV population (more heterogeneity) is, given an identical mean for the EPSP projection weights, the more tuned the PV activities to the excitatory assemblies become. This manifests itself in amplified PV responses to external sensory input. However, this comes at the cost of reducing competition between the excitatory assemblies, and therefore, less separability (and feature selectivity) between the excitatory neuronal activities. Finally, increasing input correlations can magnify these effects.

### Emergence of PV subnetworks and monotonic reciprocally tuned IPSP amplitudes generalizes to more realistic networks

To determine whether our results hold for more general and realistic cases, we considered some variations. First, we asked about the importance of tuning strength of the E networks. This was implemented by generating three excitatory assemblies with a different tuning strength, implemented by each receiving a different degree of private correlated input (Fig. 5A, c = 0.05 for E_1_, c = 0.2 for E_2_, and c = 0.4 for E_3_). Second, we introduced randomness in the E-to-PV connections. The EPSP projection weights followed a log-normal distribution (Fig. 5C), and initially there was just one population of PV cells. Finally, we were interested to see if heterogeneity in the excitatory target firing rate can affect the results. To check this, the target postsynaptic firing rates for the E cells were drawn from a log-normal distribution. As a result of these heterogeneities, the distribution of the PV cell firing rates was also log-normal (Fig. 5B).

**Fig. 5.**
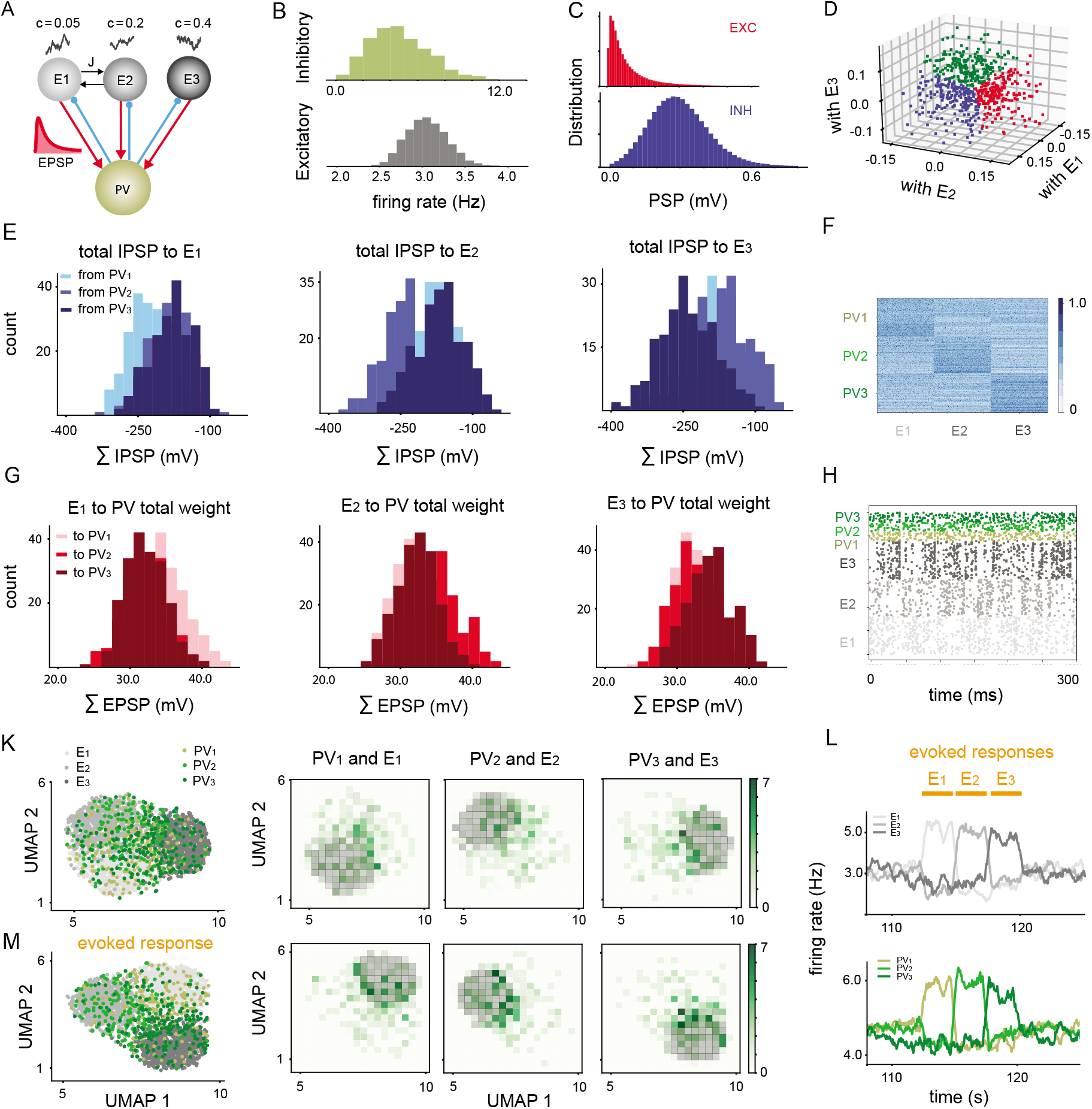
Emergence of tuned PV subnetworks in a network with heterogeneity in the input correlation as well as EPSP amplitudes and target homeostatic firing rate. **A**: Network with 3 excitatory assemblies, each receiving a different level of correlated input, and initially a single PV population. **B**: Log-normal distribution of E and PV firing rates after training for 2000 seconds. **C**: EPSP amplitudes for the connection weights from E to PV follow a log-normal distribution and are not plastic. |IPSP| amplitudes after 2000 seconds of training also follow a log-normal distribution.**D**: Response similarity between individual PV cells and each assembly population responses E_1_, E_2_, and E_3_ form a 3-dimensional cluster of vectors. After training, PV cells divide into 3 distinct groups with a slightly higher response similarity to one of the excitatory assemblies: blue: high similarity with E_1_; red: high similarity with E_2_; green: high similarity with E_3_. **E**: Distribution of the total IPSP weights from individual neurons in each PV populations to E_1_ indicate that PV_1_ has the highest preference of connection to E_1_, although these group of PV cells were labeled as PV_1_ based on their high response similarity to E_1_. Similarly, PV_2_ (PV_3_) neurons have the strongest sum projections onto E_2_ (E_3_). **F**: Connectivity matrix between labeled PV cells based on the population response similarity measure and the excitatory neurons. **G**: Total EPSP projections from individual cells in E_1_ onto the assigned PV populations indicate that E_1_ had a stronger total projection weight onto PV_1_ neurons. A similar relation holds for other excitatory assembly projections: e.g summed EPSP weights from E_3_ onto PV_3_ were more skewed towards bigger values. **H**: Raster plot of neuronal activities for the last 300 ms of training. **K,M**: 2-dimensional UMAP projection of the neuronal activities for 10 seconds of simulation time according to their distance in the high dimensional space for the spontaneous (K) and evoked (M) state. **L**: sensory input driving E_1_ (from t = 112.5 s to t = 115s post learning), E_2_ (from t = 115s to t = 117.5s post learning) and E_3_ (from t= 117.5s tot = 120 s post learning) assemblies evoke responses in PV_1_, PV_2_ and PV_3_ subnetworks, respectively.

To study the emergence of tuned inhibitory connections, initially, all IPSP projections from the single PV population onto the excitatory assemblies were chosen to be identical. 2000 seconds after introducing the plasticity rule, however, the inhibitory weights followed a log-normal distribution too (Fig. 5C). We froze the inhibitory weights and projected a sequential sensory input, targeting the E_1_ assembly first at t = 112.5 s post training, with a duration of 2.5 seconds for each excitatory assembly (Fig. 5L, top). We considered the network responses during the spontaneous and evoked state from t = 110 s to t = 120 s, for the rest of the analysis. Each PV cell projected to E cells in 3 different assemblies, characterized by their independent and private source of correlated external input. Therefore, for each PV cell, one can derive a 3 dimensional vector with each element representing the response similarity between the PV cell and the population responses from the 3 distinct excitatory assemblies (Fig. 5D). While the responses clearly do not form distinct clusters, we can nonetheless assign PV neurons a group identity according to the maximum absolute value of the components of each vector; i.e a PV cell with a maximum response similarity to the population response of E_1_ was labeled as PV_1_. This way, we separate the cloud of vectors into three distinct PV populations (Fig. 5D). We call this measure of labeling *population similarity measure.* We used 3 other measures, namely the *individual similarity measure*, where the average response similarity between *pairs* of PV and E cells were used to label PV cells; the *outgoing PV measure* where the sum of the outgoing connections from each PV cell to the neurons in the 3 excitatory assemblies were measured and the maximum projection weight among the 3 vector components defined the label for individual PV cells (Fig. A4 and A5); and the *incoming E measure* which calculates the total EPSP projection weights onto each PV cell and the label of the PV cell is defined based on the maximum incoming weight from each E assembly. The intersection of PV labels for all these 4 measures are shown in Fig. A4C, D, indicating that individual and population similarity measures result in the highest overlap between PV labels, but in general, all measures have a significant overlap between the labels.

To determine whether using the population similarity measure to assign labels to PV cells revealed any specific pattern of connectivity between the E and PV cells, we plotted the distributions of the total IPSP weights from individual PV cells to E_1_ through E_3_ (Fig. 5E). We found that PV_1_ projected strongly onto E_1_ compared to E_2_ and E_3_. This result indicates that clustering PV cells based on their response similarities to the population signals from the incoming E assemblies can result in PV labels that reflect a structured connectivity matrix (Fig. 5F).

The underlying cause for the clustering of PV cells was random stronger excitatory drive from distinct excitatory assemblies to a portion of the PV cells, as shown in the distribution of the total excitatory weights from E_1_ through E_3_ to individual PV populations (Fig. 5G). In other words, E_1_ had the strongest projection weights onto the group which we characterized as PV_1_ (Fig. 5G, left). Similarly, E_2_ and E_3_ had the biggest total EPSP weight to PV_2_ and PV_3_, respectively (Fig. 5G, middle and right). This is consistent with the understanding provided by our mean field theory in Fig. 2.

How do neuronal activities in the spontaneous state relate to the evoked state? We observed that the raster plots of neuronal activities in the spontaneous state showed weak correlations in activity between PV and E cells (Fig. 5H). To evaluate feature tuning in this more realistic network example, we first plotted the neuronal activities within the last 7.5 seconds of spontaneous activity in a two-dimensional UMAP distance space (Fig. 5K). Excitatory neurons within each assembly formed distinct clusters with slightly intermixed borders. The PV cells clustered in the middle of the E clusters, demonstrating low similarity with the E cells. We also plotted the distribution of the neurons in the 2D UMAP plane (Fig. 5K). The degree of overlap between the distributions of PV and E neurons on the 2D UMAP plane reflects the degree of co-activity between the cells.

Evoked responses were obtained from network simulations in which the external input to individual E populations were increased by %3 (Fig. 5L). This resulted in increased covariance (and hence response similarity) between co-tuned *E* and *PV* cells, and as a consequence, the overlap between the distributions of their reduced responses in the 2D UMAP plane increased. Also, PV subnetwork activities formed a denser cluster around the E assemblies to which they were co-tuned but the distribution of the PV cell responses on the 2D plane was still broader than those of the E cells, which indicates broad tuning of PV cells in general. This finding is consistent with most of the experimental reports [1–3]. We conclude that the activity in the spontaneous state could also reflect feature tuning of PV cells in the evoked state.

The results of this section indicate that for more realistic networks with heterogeneities in the target excitatory firing rates, EPSP amplitudes and levels of shared correlated input, tuned responses in PV cells emerge according to the distribution of the EPSP projecting weight, and correlations play a role in shaping the tuned PV weights. We showed that sensory input to individual excitatory assemblies can drive the activity of the corresponding PV cluster, and this indicates co-tuning of PV cells at a population level. However, the broader distribution of individual PV cell responses indicates broad tuning of individual PV cells.

### Co-tuning of PV cells is a network property which is not reflected in pairwise correlations

Similar to the experimental reports in layer 2/3 of mouse visual cortex [6, 17], we observed a monotonic relation between the EPSP and IPSP amplitudes in all reciprocal motifs in the network under study (4000 samples of data were plotted in Fig. 6B left). This generalizes our earlier results in Fig. 4 to a more realistic scenario since for reciprocally connected E and PV pairs, an increased EPSP increases the correlation (and hence response similarity) between the pairs. As a result of the symmetric plasticity curve, the IPSP amplitude increases with correlation (or equivalently, response similarity Fig. 6A). However, if the EPSP values become too strong, while the correlation between E and PV pairs increases, the IPSP values start to decrease (this becomes apparent when all samples are plotted, but since such cases comprise only a small percentage of all PSP values, we did not observe them in the 4000 samples of the PV-E pairs in Fig. 6B). This occurs because the increased firing rate of the PV cell balanced the firing rates of the E cells by reducing the projecting IPSP value (See Fig. A3). This suggests that very strong EPSP values can potentially create lateral inhibition within each excitatory assembly.

**Fig. 6.**
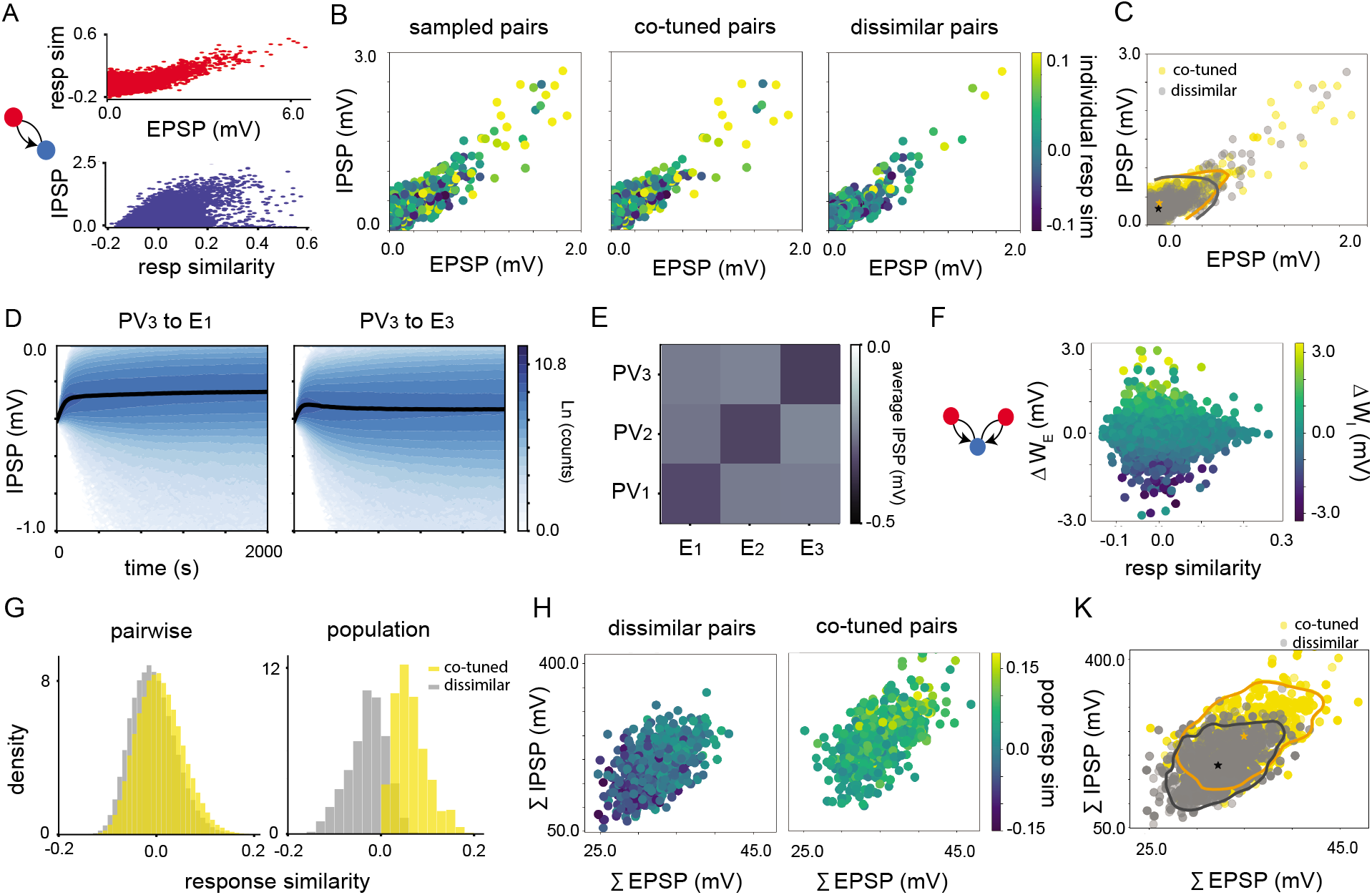
PV tuning is a network property. **A**: For reciprocally connected E and PV cells in the evoked state, response similarity between the pairs increases with increasing values of EPSP, and IPSP increases monotonically with pairwise similarity. **B**: Left: EPSP and IPSP values for 4000 sample of reciprocally connected pairs. Middle and right: Reciprocal EPSP and IPSP values for pairs belonging to similarly tuned populations and dissimilarly tuned populations. Colors represent pairwise response similarity between the E and PV cells. **C**: Scatter plots of co-tuned and dissimilarly tuned PSP pairs overlayed on one another and the contour lines indicate regions which contain at least %95 of the data from both categories. **D**: Evolution of IPSP distributions from PV_3_ cells to E_1_ (dissimilarly tuned) and E_3_ (similarly tuned) cells as a function of time. **E**: Average connectivity matrix for the weights from PV subnetworks onto excitatory assemblies. **F**: Difference between the IPSPs for the triplets of E-PV-E as a function of response similarity between the excitatory cells and the difference between the projecting excitatory weights onto the shared PV cell. As Δ *W_E_* increases, Δ *W_I_* also increases. **G**: Distribution of pairwise response similarities between E and PV cells for co-tuned and dissimilarly tuned pairs are almost identical (left), while distributions of the response similarities between individual PV cells and the population signals from individual E assemblies are more significantly different for co-tuned and dissimilarly tuned pairs. **H**: Total IPSP projection from individual PV cells onto similarly (right) and dissimilarly (left) tuned E cells as a function of the sum of the EPSP received by the E assemblies. The colorbar defines similarity of the PV cells with the average excitatory populations **K**: Distribution of summed IPSP as a function of summed EPSP for co-tuned (yellow) and dissimilarly tuned (gray) PV cells to the E assemblies. Stars indicate the mean of the distributions, and boundaries define regions in the space which contain %95 of the data from each group.

It is tempting to think that E-PV pairs with strong reciprocal weights and high correlation coefficients (response similarity) belong to similarly tuned assemblies, while those with weak correlation coefficients and weak mutual synapses belong to dissimilarly tuned subnetworks. To check this, we plotted 2000 samples of pairs which belonged to co-tuned subnetworks (Fig. 6B, middle) and 2000 samples which were not co-tuned but belonged to dissimilarly tuned subnetworks (Figure 6B, right). Interestingly, we observed that there was almost no distinction between the strength of the synaptic patterns (Fig. 6C) or their pairwise response similarities (correlation coefficients). In fact, the distributions of IPSPs for the PV cell projections to co-tuned or dissimilarly tuned E neurons quickly evolve toward very high variances (Fig. 6D, the variance of IPSP amplitude for the connections with similarly tuned E cells is slightly bigger). This high variance accounts for the observations in Fig. 6B which show almost no distinction between the ranges of EPSP and IPSP values for co-tuned and dissimilarly tuned pairs. However, the mean of the distributions reflect the tuning of the subnetworks. The average connectivity matrix from PV to E assemblies has a diagonal structure which reflects stronger tuned weights from the PV populations to their corresponding co-tuned E assemblies (Fig. 6E). The strongest correlated external input drove the establishment of the strongest connectivity, here from PV_3_ to E_3_ (*c* = 0.4), relative to that of E_2_ and E_1_.

We were also interested to check if under these plasticity rules, the connections from a PV cell differentiate between two E cells with high or low response similarity (or equivalently correlation coefficient between the E cells). Due to shared correlated input, the response similarity between E cells in recordings under natural input conditions should give a hint as to whether the two E cells belong to the same assembly or not. We assumed that we did not know the source of the correlated input and only relied on the pairwise response similarity between pairs of E cells which had one projecting PV cell in common, and searched for all triplets like the cartoon in Fig. 6F. We plotted the similarity between the E cell responses on the horizontal axis and the difference between the projecting EPSPs from the E cells onto the PV cell (Δ*W_E_*) on the vertical axis. Then, the difference between the reciprocal projecting IPSP from the PV cell onto the E cells were plotted in color codes (Δ*W_I_*). As expected, due to the monotonic relationship between EPSP and IPSPs in the network, for increasing values of Δ*W_E_*, we observe an increased value of Δ*W_I_*, which shows the important role of heterogeneity in shaping PV weights. However, we observed that for a fixed value of Δ*W_E_*, as the response similarity (correlation) between the two E cells increased, the difference between the projecting IPSP amplitudes Δ*W_I_* did not change. This highlights the fact that pairwise correlations between two E cells and between PV and E cells do not reflect tuning of the cells, despite the monotonic relationship between Δ*W_E_* and Δ*W_I_*.

In fact, pairwise measures such as response similarity between a pair of connected PV and E cell cannot give a significant indication about the membership of individual PV cells in co-tuned or dissimilarly tuned subnetworks. The distribution of the response similarities between pairs belonging to any of those groups is almost identical (Fig. 6G, left) and they are not significantly different from one another. However, if population measures (such as “population similarity measure”) are used to determine the tuning of PV cells, then the distribution of the co-tuned versus dissimilarly tuned PV cells to E assemblies are significantly different (Fig. 6G, right). Utilizing other network-level measures, such as “outgoing PV measure” (Fig. A5) or “incoming E measure”, also results in more significant separation between the distributions of co-tuned and dissimilarly tuned PV cells.

Further, if the total sum of projecting EPSPs from E assemblies to single PV cells (and total IPSP projections from single PV cells onto the E assemblies) are taken into account, then co-tuned and dissimilarly tuned PV cells can be better identified (Fig. 6H). Higher values of the summed EPSPs would, on average, result in higher summed IPSP values and higher population response similarities (Fig. 6H, co-tuned pairs). If PV cell connections are classified into two binary groups of co-tuned and dissimilarly tuned connections, the dissimilar connections will be associated with small total EPSP projections from the corresponding E cells which results in smaller summed IPSP values, and co-tuned connections will cluster in the region with high values of summed EPSPs with higher summed IPSP values (2000 samples from each case plotted in Fig. 6K). This population-level measure can create more separable and identifiable distributions in terms of weight tuning for co-tuned and dissimilarly tuned cells. This is more easily seen when the “outgoing PV measure” is used to classify the PV cells (Fig. A5 H, K).

These results demonstrate that for a more realistic network with heterogeneities, our theoretical results for simpler networks still hold, namely, there is a monotonic relation between the reciprocal EPSP and IPSP amplitudes for pairs of connected E and PV cells. However, high values of pairwise response similarity do not necessarily imply that the two cells belong to co-tuned subnetworks of PV and E cells. Instead of a single EPSP projection value from the presynaptic E cell (individual measure), if the total sum of the projecting EPSPs from different E assemblies (population measure) are considered, then PV cells which receive a large value of this quantity would be co-tuned to the corresponding projecting E assembly and the similarity between the PV cell and the E population signal would take higher values. This population measure provides a more reliable quantity to determine the tuning properties of individual PV cells, as the distribution of the similarities are more distinct.

### Co-tuned E and I subnetworks exhibit more stability and a wider dynamical range of network responses

One advantage of a network with multiple excitatory and inhibitory populations with tuned connections is that in these networks excitatory neurons become more decorrelated from each other [25] and as a result they can maintain stability [6] for a wider range of parameter changes in the circuit. To investigate the response dynamics and stability of the networks under different distributions of E to PV cells, we chose the two networks with fixed in-degree and log-normal E to PV distributions in Fig. 1. We froze all plastic weights at t=2000 s, and compared the raster plots, covariance functions, power spectral densities, and the eigenvalue spectrum of the networks for *w* = 2.5, i.e, the scalar defining the strength of the excitatory connections within each assembly.

Activity in the network with fixed in-degree distribution (Fig. 7A) expresses stronger competition between the excitatory assemblies (Fig. 7B) with a relatively longer time of active state for each assembly (Fig. 7A). However, the network with a log-normal distribution shows more rapid switches between the high and low active states for each assembly dynamics (Fig. 7A) with relatively shorter active state for each assembly, and also weaker competition between the excitatory assemblies (Fig. 7B). The normalized power spectrum for the average neuronal dynamics in E_1_ (similarly for E_2_) indicates stronger high frequency responses for the network with log-normal distribution. The relative high frequency power for E dynamics in the network with fixed in-degree is almost one order of magnitude less (Fig. 7C).

**Fig. 7.**
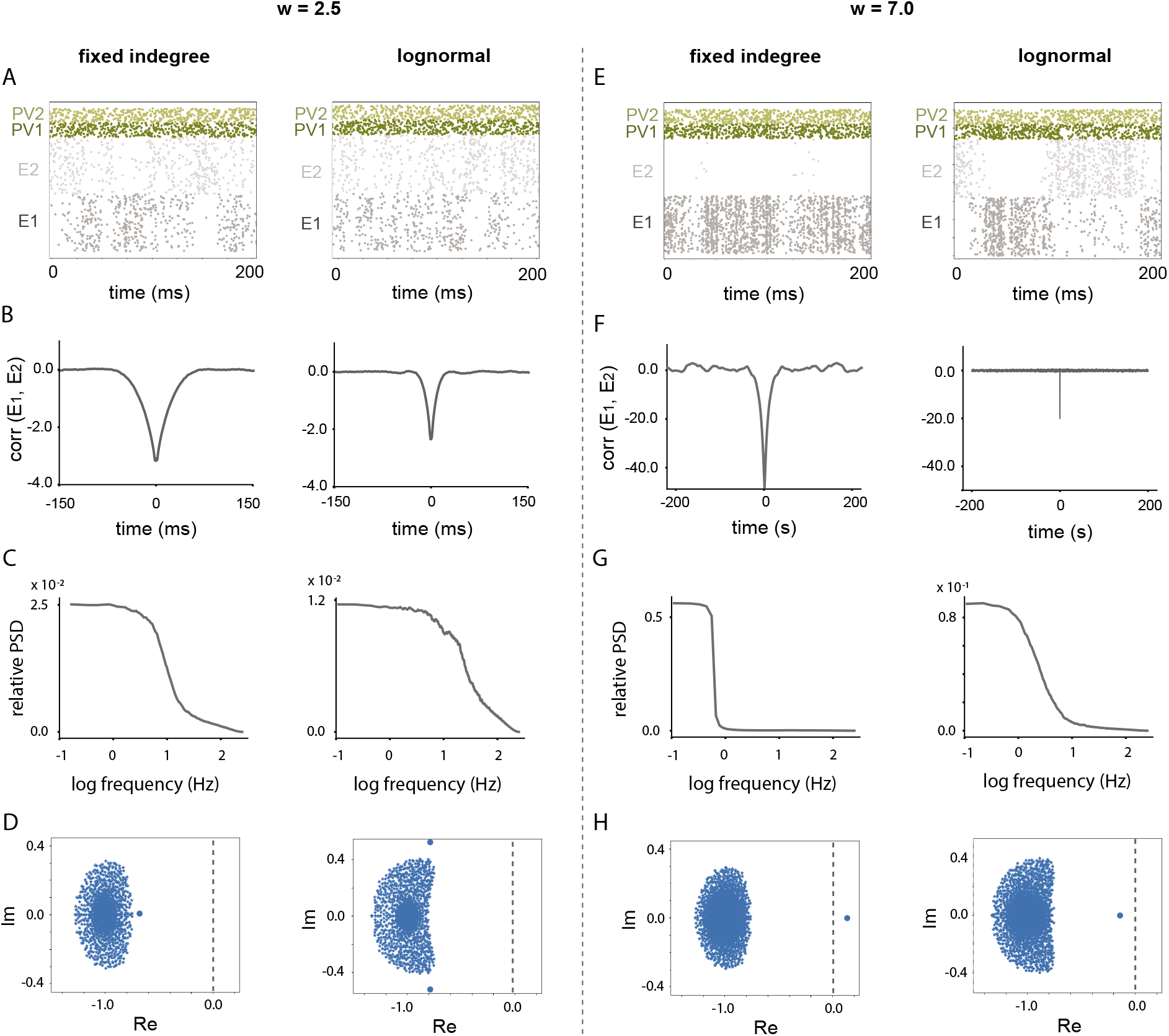
Effect of increasing E to E weights within assemblies on network stability and frequency response range. **A**: Raster plots of the network activity for *w* = 2.5, for the networks with fixed in-degree (with one population of PV cells which did not split) and log-normal E to PV weight distributions (with self-organized two populations of PV cells with reciprocal connections), respectively, with frozen weights obtained after 2000 seconds of training in Fig. 1. **B**: Cross-covariance function between the average population activities of the excitatory assemblies. **C**: Power spectral density of the E_1_ firing rate for the fixed indegree and log-normal network, respectively. **D**: E_1_genvalue spectra of the linearized neuronal dynamics for networks in A and B, respectively. All eigenvalues have negative real parts. **E**: Raster plots for stronger within assembly connections (*w* = 7.0) for networks in A and B, respectively. **F**: Cross-covariance function between E_1_ and E_2_ average neuronal activities. **G**: Power specral density for the firing rate of E_1_, in the network with fixed in-degree and log-normal distribution, respectively. **H**: Eigenvalue spectrum of the network dynamics for (*w* = 7.0) for networks in K and L, respectively. The network with only one PV population has one positive eigenvalue (and is unstable); however, the network with co-tuned E and PV subnetworks has remained stable.

The eigenvalue spectrum of the network with fixed in-degree distribution and that of the network with log-normal distribution (Fig. 7D, taken from the Jacobian of the high dimensional system) shows a denser inner bulk within a bigger cloud of eigenvalues.Since all eigenvalues are negative, both systems are stable. Further, as the absolute value of the leading eigenvalue for the network with fixed in-degree is smaller than that of the network with log-normal distribution, the switching dynamics between the excitatory assemblies is slower (with longer active time for each assembly) for the network with fixed in-degree distribution. Note that in both cases there are two additional complex conjugate eigenvalues with absolute values of their real parts bigger than 2.5, but they are not shown in Fig. 7D,H.

To test the stability of these networks under parameter changes, we increased the value of *w*, and strengthened the connections within each excitatory assembly (*w* = 7.0 here). However, we did not scale the connections from PV to E cells. As a consequence, the raster plots (Fig. 7E for the fixed in-degree network and for the log-normal network) showed stronger degrees of synchrony between the activities of the excitatory cells within each assembly (more synchrony for the network with fixed in-degree). Also, there was a much stronger level of competition between the activities of the two assemblies for the network with fixed in-degree distribution (Fig. 7F).

Increasing the weights within assemblies had a drastic effect on the switching dynamics of the E assemblies and made the dynamics much slower compared to the case with *w* = 2.5 (Fig. 7E, G). Here, again, the active state for E_1_ in the network with fixed in-degree was much longer than that of network with log-normal distribution. This can be clearly seen in the narrow power spectrum of the average firing rate of E_1_ for the network with fixed in-degree as compared to the wider power range for the dynamics of E cells for the network with log-normal distribution (Fig. 7G). This example shows the role of reciprocal PV connections in increasing the frequency range of the excitatory neuronal dynamics.

For both networks, the eigenvalue spectra associated with each network for *w* = 7.0 had a leading eigenvalue well separated from the bulk of the eigenvalues. For the network with fixed in-degree, this eigenvalue has already crossed the boundary line (edge of stability) at zero (Fig. 7H), and represents an unstable system. This instability manifests itself in the extremely long active state of the excitatory populations as well as the high level of synchrony between the excitatory cells (due to firing rate homeostasis with fast inhibitory time scale, the mean of the activities cannot go beyond the target value, and instability manifests itself in the higher level of synchrony). However, all eigenvalues of the network with log-normal distribution, including the leading eigenvalue, lie on the left side of the stability boundary, and therefore, the network remains stable, even for strong connections within each excitatory assembly (Fig. 7H).

These results indicate that if due to learning, the connection strengths within excitatory assemblies increase, then the network with a high level of heterogeneity in connections from E to PV, which receives reciprocal connections with proportional strengths back from PV to E cells, will be better able to maintain the stability of the network with less pathological E activity (less degree of synchrony). Also, we showed that reciprocal connections from PV cells to E cells that respect the projections from E to PV cells proportionally, result in an expanded frequency range with relatively stronger components for the excitatory responses.

## Discussion

In this study, we showed how PV cells can develop co-tuning with E cells by considering the “network effects” of heterogeneity in the EPSP connections from E to PV cells and correlated shared input from other layers. More specifically, we used a known homeostatic inhibitory plasticity rule to show the self-organization of initially untuned PV cells into subnetworks with relatively high correlations with the excitatory assemblies. This suggests that the pooling hypothesis [1], when supplemented with correlation patterns of excitatory assemblies and the heterogeneity of the EPSP amplitude, can capture the co-tuning of PV with E cells, reported in [5, 6], as well as the monotonic relationship between EPSP and IPSP in reciprocally connected E and PV cells [6, 17].

We reconciled conflicting studies on the tuning responses of PV cells by highlighting the difference between individual and population-level measures for tuning. More specifically, for a network simulation with heterogeneous parameters, highly correlated pairs with strong reciprocal EPSP and IPSP were found in both co-tuned and dissimilarly tuned E and PV assemblies. Therefore, pairwise response similarity between E and PV cells may not provide a good measure to determine the tuning of the PV cells. Pairwise measures result in distributions of response similarities (or equivalently, correlation coefficients) for cotuned and dissimilarly tuned pairs that are not significantly different from each other. In contrast, population measures such as population response similarity result in well-separated distributions for co-tuned and dissimilarly tuned PV cells.

A fact that plays a crucial role in our study is the diversity and heterogeneity of EPSP amplitudes from E to PV cells. Intracellular *in vitro* recordings reveal that in layer 2/3, the EPSP amplitudes from E to PV cells are much stronger and more diverse than the connections between E cells [2, 14, 21]. Moreover, highly feature-tuned PV cells have smaller dendritic fields, but broadly tuned PV cells have longer dendrites with larger dendritic fields [29]. This suggests that the overall excitatory input to PV cells shapes their tuning. Consistent with this, we showed that this high variance of the connection strength from E to PV cells can by chance correlate some groups of PV cells to some excitatory assemblies. This would create high response similarity between E and PV cells [4, 6] and results in co-tuning of PV cells to a group of E cells. This is consistent with the pooling idea but, most importantly, includes the effect of EPSP variance and heterogeneity into account.

Another component that we employed in our modeling was a source of shared correlated input. We assumed that this external source defines the tuning of excitatory neurons within an assembly to different input features. It is known that shared input from cortical layer 4 and also within layer 2/3 forms fine-scale subnetworks of excitatory neurons in layer 2/3 [19] with stronger connections between them [30]; however, the connection strength between uncorrelated E cells become weaker. These strong E to E connections play a role in shaping stimulus-specific feature selectivity [20]. In mouse V1, it was shown that the pattern of correlation between excitatory neurons is strongly influenced by the external stimuli, emphasizing the importance of feedforward connections in shaping responses among E cells. However, correlation patterns among PV cells are much less affected by the external input but rather are shaped by the local projections from E cells within layer 2/3, reflecting the importance of recurrent connections in driving response patterns of PV cells [2]. We employ this evidence in our computational study and show that excitatory assemblies formed via shared correlated input can drive the responses of PV cells through recurrent connections and form co-tuned subnetworks.

The last component that we considered in our modeling was a homeostatic plasticity rule that governed the connections between PV and E cells. It is known that in layer 2/3 of the visual cortex, when excitatory neurons have a bursting activity, long-term depression (LTD) is induced if the presynaptic fast-spiking interneuron fires within 100 ms after the excitatory burst [26]. Also, in the mouse auditory cortex, a symmetric STDP curve with potentiation of the inhibitory synapse for spike timing of E and I cells within a short time frame of 10 ms governs the synaptic changes [23]. In hippocampal slices, however, a longer interval between the spike emission of E and I cells results in the suppression of the synapse [22]. A similar STDP curve was also recently reported to describe the inhibitory synapses from PV cells onto E cells in the orbitofrontal cortex of mice [25]. This kind of STDP has been shown to result in postsynaptic firing rate homeostasis [24]. Based on this evidence for the activity dependent dynamics of synapses from GABAergic PV interneuron to E cells, we employed a similar symmetric STDP curve for those connections in our study. Given the slow time scale of inhibitory plasticity [31], we assumed that the excitatory assemblies were already formed through shared correlated input from layer 4 (Fig. A1).

Correlations between E and PV cells played an essential role in shaping the tuned connections from PV to E cells and creating a monotonic EPSP and IPSP for the reciprocal connections. Ignoring the correlation between E and PV populations does not capture the tuned weights observed in the simulations. In a recent study [32], this correlation term was ignored, and the authors did not capture this phenomenon by only considering heterogeneity in excitatory weights. In [32], plastic excitatory weights for the connections from E to PV cells had to be considered, and even a correlated growth dynamics for the reciprocal EPSP and IPSP amplitudes had to be imposed (embedding the observed monotonicity in [6] between EPSP and IPSP in the weight update instead of this relationship emerging from the dynamics). Since the authors used a stimulus specific form of homeostasis to embed assemblies, inhibitory assemblies were formed. However, here, we did not assume any correlations in the weight updates but showed that the correlations between the activities of the E and PV cells would be sufficient to get reciprocal and monotonic EPSP and IPSP weights, consistent with experimental findings [33]. It is important to note that in our simulations, for very strong EPSP amplitudes, which may result from a very wide distribution of EPSP onto PV cells, the reciprocal IPSP amplitude decreased. Such cases comprised a very small percentage of our data, which did not reflect in our samples in Fig. 6. This suggests that within similarly tuned subnetworks, E-PV-E motifs within the same or different E assembly may exist, and this can cause lateral inhibition with emerging competition properties. However, in our simulations, the relationship between EPSP and IPSP was monotonic for the majority of the synapses. Considering the spatial dependence of EPSP amplitudes [6, 34, 35], this emerging property provides a detailed balance between excitatory and inhibitory neurons [24], therefore, neurons are maintained in the balanced state over time [36–39], and over space [17]. This can be important in stabilizing spatiotemporal patterns in the cortex.

There is a direct relationship between competition between E assemblies in a network and its feature tuning and input amplification. In our study, sensory input to one of the excitatory assemblies caused a significant input amplification for the delta network (no variance in the E to PV connection strengths) but no differentiation in PV responses. However, PV responses changed in the wide network in response to sensory input to one of the excitatory cells. Feature selectivity, input amplification, and competition are all compromised in wide networks due to the tight tracking of excitatory activities by the PV subnetworks. In light of the balanced amplification idea [40], the difference between the excitatory and inhibitory firing rate fluctuations in co-tuned subnetworks of the wide network are smaller than those in the delta network. Therefore amplification of the external input (and equivalently, fluctuations in the spontaneous state) for the wide network will be weaker. However, for the delta network, the inhibitory firing rate is almost constant, providing global balance and keeping a relatively fixed sum of all the excitatory firing rates in all assemblies combined, with relatively little fluctuation in the inhibitory subnetwork [41]. This can result into an attractor network which is mediated by uniform inhibition [41, 42].

In our study, we did not consider somatostatin (SOM) inhibitory subtypes. It was shown that in OFC, SOM cells expressed Hebbian plasticity in their connections to to E cells [25]. If SOM cells are tuned to a specific excitatory population [43], lateral inhibition between E assemblies emerges [25]. This study showed that random projections from SOM to PV cells can by chance target a group of PV cells more strongly than other groups. This causes correlations between those PV and E cells that are strongly impinged upon by the SOM group. As a result of the high correlation between those PV and E cells, mediated through the SOM group, strong connections from those PV cells to the E assembly will emerge. It would be interesting to study the role of random projections and heterogeneity from both E to PV and SOM to PV cells and investigate if any competition between SOM and E cells in determining the pattern of PV connectivity emerges. Additionally, SOM cells participate in a disinhibitory circuit through the connections from VIP to E cells and may paint a more complicated picture when considered in this analysis [44].

We showed that reduced selectivity and input amplification in reciprocally connected E and PV subnetworks comes as costs for providing circuit stability [6], as shown in the eigenvalue spectrum, and increased dynamical range of the excitatory cells. In control theory, negative feedback can expand the system’s dynamic range and provide robustness to noise. Previous theoretical results also show that the inhibitory structure that emerged in our study provides a broader frequency range for network operation [45]. Since the reciprocal PV to E connectivity structure expands the frequency response of the spontaneous activity, it can also play a role in shaping gamma oscillations. In fact, it was observed that inhibiting PV cells caused a suppression of the gamma oscillations *in vivo* [46, 47], and their activation caused gamma oscillations [46]. Both gamma oscillations [46] and strong reciprocal excitatory to inhibitory connections [48] have been shown to enhance signal propagation and reduce noise in the network. The theory developed in our study predicts that the reciprocal connections are responsible for this noise attenuation.

## Methods

### Network models

Network simulations were conducted in NEST [49], using Leaky Integrate-and-Fire (LIF) neuronal dynamics with current based synapses with delta functions. In all network simulations, PV neurons composed %20 of the network size and the rest of the network was composed of excitatory cells. In our study, we considered different networks with different sizes and parameters. More detailed information is available in Appendix A.

For all simulations, the dynamics of each neuron follows

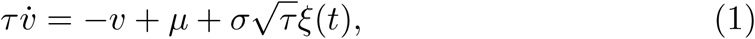

in which *μ* and *σ* are the mean and standard deviations of the total (external plus recurrent) input to the cell. In all network studies in the main paper, only the synapses from PV to E cells were plastic and followed a symmetric STDP function [24] as follows:

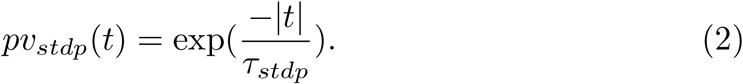

The variable *t* represents the time interval between the pre and postsynaptic spikes, and as opposed to Hebbian learning, the causal relation between them does not affect the synaptic weight update. Spike emission by the presynaptic PV cell causes a depression of the synapse. These conditions were shown to be required for the plasticity rule to ensure postsynaptic firing rate homeostasis [24].

The weight dynamics from PV to E cells employing the homeostatic plasticity rule in [24] are composed of two parts, as follows:

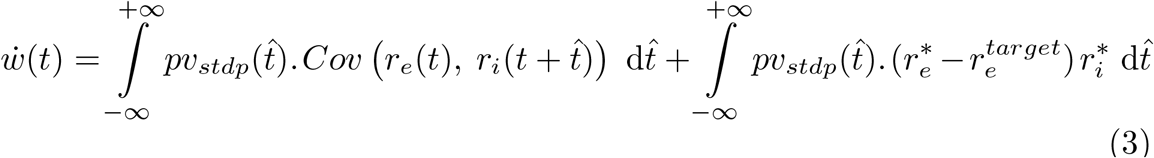

The first term on the right hand side of the equation above reflects the contribution of the cross-covariance function between the pre and postsynaptic cell activities, represented by 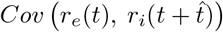. We call this term the “**learning signal**” which we utilized to justify growth and decay of some inhibitory weights, specially after firing rate convergence. The second term represents the effect of mean firing rates (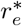 for the excitatory postsynaptic neuron and 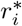 for the inhibitory presynaptic PV cell). Transient firing rates converged to stationary solutions within the first few ten seconds of the simulations, in most of our network studies. However, given a constant stationary background input, most of PV to E weight changes happened after this settlement to the stationary mean solution. A mean-field analysis to explain the emergence of PV weight tuning on the network level can be found in the Supporting Information in Appendix A.

We considered E to E Hebbian plasticity, as well as E to PV Hebbian plasticity only in Fig. A1 to show that our results do not change when excitatory weights also obey a standard plasticity rule, such as Hebbian STDP.

### Data analysis

To calculate the response similarity between PV and E cells, we considered individual instantaneous neuronal firing rates as vectors in a high dimensional space and calculated the cosine between them. In other words, we used the following formula:

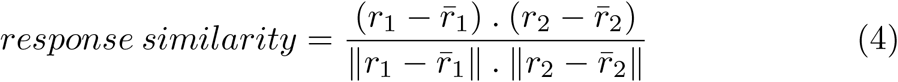

where *r*_1_ and *r*_2_ are the firing rates of the presynaptic and postsynaptic cells, and ||·|| is the Euclidean norm of the responses for each cell.

To reduce the high dimensional activities of the cells, we used Uniform Manifold Approximation and Projection (UMAP) for dimension reduction [28], and we used the *cosine* measure to identify the distance between data points for the reduced dimensional representation. This method uses algebraic topology to preserves more of the global structure of the data, compared to any other dimension reduction methods.

Average neuronal covariance functions in Fig. 2 were obtained from average instantaneous neuronal firing rates in *E*_1_, *E*_2_ and *PV*_1_, in temporal resolution of 1 ms. Fluctuations in the neuronal activities of the network for an interval of 10 seconds before and after synaptic updates were projected onto a 2D UMAP space (average neuronal responses were subtracted from the instantaneous firing rates).

To investigate the role of heterogeneity in connection strengths from E to PV (and hence from PV to E cells as a result of plasticity) in shaping the tuning responses of the E cells in Fig. 4, we presented a specific sensory input (with shared correlated and common uncorrelated components) to the cells in E_1_. This sensory input was also composed of shared and common components which affected all the excitatory cells in the network. This stimulus was applied to the network with frozen weights (obtained after 1000 seconds of simulations with STDP) at t = 120 s, and continued for 50 seconds. Average neuronal firing rates for the E and PV populations were plotted for a 100 second interval which included this sensory input (t = 100 s to t = 200 s, Fig. 4L). UMAP plots were obtained from mean-subtracted neuronal responses from t = 115 s to t = 125 s (Fig. 4H).

For the network simulations in Fig. 5 and Fig. 6, small sensory inputs (%2 of the total background input to the E cells) were presented to E_1_, E_2_, and *E_3_* populations at t = 112.5 s, t = 115 s and t = 117.5 s, respectively, with frozen weights obtained 2000 second after the onset of the inhibitory STDP rule. Labeling of the PV cells was based on the PV responses to these external sensory inputs, and their response similarity with the excitatory population responses (population response similarity measure). Instantaneous firing rates of all neurons were calculated in temporal resolutions of 15 ms, for a total duration of 7.5 seconds (from t = 112.5 s to t = 120 s). In order to show the response similarity of the PV and E cells, we subtracted the mean of the neuronal responses. These neuronal fluctuations were used for the UMAP plots in Fig. 5.

## Acknowledgments

FL and ALF thank Petr Znamenskiy, Eric Shea-Brown, and Hannah Bos for helpful discussions and comments on this version of the paper. FL is supported by the Swartz fellowship in Theoretical Neuroscience. This project was also supported by the Simons Collaboration for the GLobal Brain (to ALF).

## Appendix A Supporting Information

### Theoretical results for the emergence of PV weight tuning

We study the interactions between the dynamics of leaky integrate-and-fire (LIF) neurons and the evolution of plastic weights between neurons in different populations. We show how the inhibitory weights self organize to form stronger connections to those excitatory populations which project onto them with a stronger excitatory weight.

The membrane potential dynamics for an LIF neuron which is isolated from the network is a low pass filter of first order with the dynamics

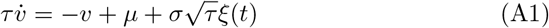

where *μ* and *σ* are the mean and standard deviation of the input signal and *ξ*(*t*) is assumed to be a zero-mean normal Gaussian white noise. When this neuron is embedded in a network, the mean and standard deviation of the total input (external plus recurrent input) have to be calculated self-consistently [50].

What correlates excitatory cells is a source of shared input which projects identical spike trains to all excitatory neurons with equal weight. While inhibitory neurons receive independent background Poisson input with a standard deviation of 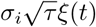 affecting their membrane potential, excitatory neurons receive inputs from two sources: a shared Poisson source which is common between E cells within an assembly and has a standard deviation equal to 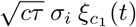, and a background independent source with a standard deviation of 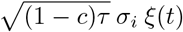. The total variance of the entire external input impinging on E cells is equal to that of received by the PV cells.

In order to get neuronal correlation functions, it is required to linearize the dynamics of individual neurons around their operating stationary firing rates. A transfer function that relates the stationary input to the firing rate of the neuron is obtained from the stationary solution of the Fokker-Planck equation which solves a first passage time problem. The stationary firing rate *r* is obtained from *r* = *f* (*μ,σ*) where the transfer function *f* (.) follows [50]

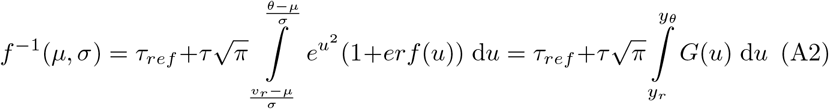

To linearize the transfer function around the operating point *r*\*, we take the derivative of both sides with respect to *μ*. This results in

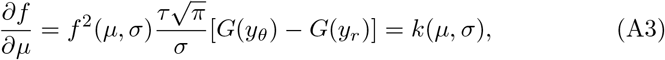

where *k*(*μ, σ*) represents the slope of the *f* – *I* curve at the linearization point, and 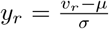 and 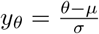.

We choose *E*_1_ and *E*_2_ to represent the average fluctuations of the firing rates of individual neurons in the excitatory assemblies, and *I*_1_ and *I*_2_ to stand for the dynamic firing rates of the PV neurons. In our model, a stronger connection weight from *E*_1_ to *I*_1_ compared to *I*_2_ differentiates *I*_1_ from *I*_2_. There exists a similar stronger connection weight from *E*_2_ onto *I*_2_ neurons.

The operating point for excitatory and inhibitory neurons are 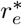 and 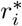, respectively (the stationary solution comes from the solution of coupled Fokker-Planck equations). Each excitatory population is composed of *N_e_* neurons, and each inhibitory population has *N_i_* neurons. The probability of connection between excitatory neurons is *p* = 0.1, and every other probability of connection is 4*p* = 0.4 (hence the factors 4 in equation A4). The connection weights from an excitatory population *j* to an excitatory population *i* are *w_ij_*. Two excitatory neurons residing in different excitatory assemblies, if connected, have a PSP amplitude of *J mV*. However, neurons within each excitatory assembly are more strongly connected (PSP amplitude of *w J,* where *w* > 1). Projection weights from excitatory onto co-tuned inhibitory neurons are equal to *q J*, with *q* > 1, however, projection weights from excitatory to dissimilarly tuned inhibitory neurons are equal to *J*. The variable *w_ik_* represents the average connection weight from a PV neuron onto an excitatory neuron in assembly *k*. The fixed (non-plastic) inhibitory weights between inhibitory neurons are –*gJ*, where *g* = 10. All inhibitory weights from *I*_1_ and *I*_2_ to all excitatory neurons are initially equal to –*gJ,* however, inhibitory plasticity will change these weights, and we are interested in dynamics of those changes.

The average dynamic mean-field equations for individual neurons around their steady state values are

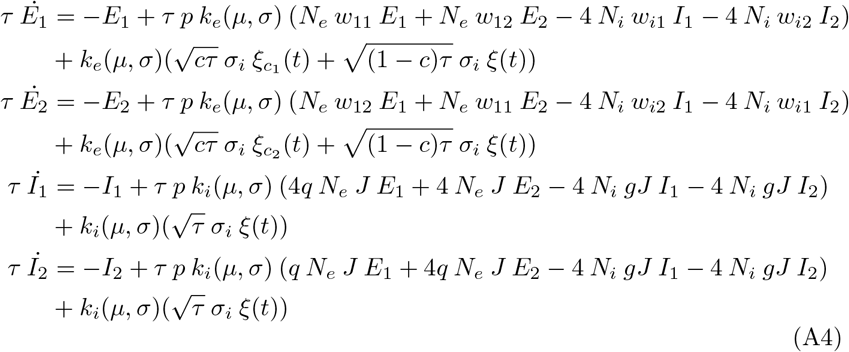

which in matrix form can be represented as

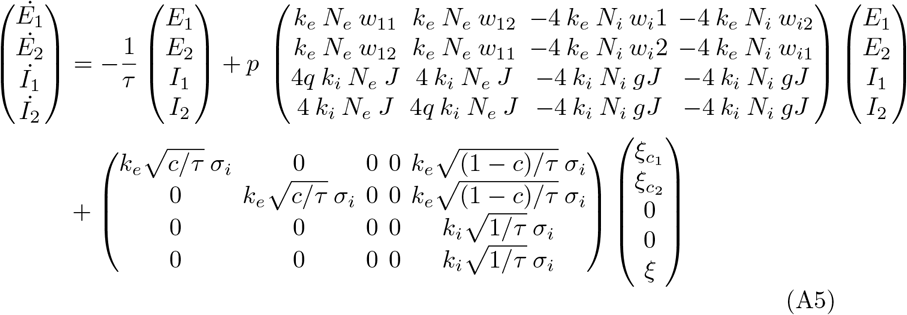

Note that *σ_i_* which appears as the standard deviation of the external inputs is different than *σ* which appears in the slope of the f-I curve. The latter has two components which include the external standard deviation and also the internally generated standard deviation [50]. The fluctuation of the rate dynamics around the fixed point can be written in the following general form:

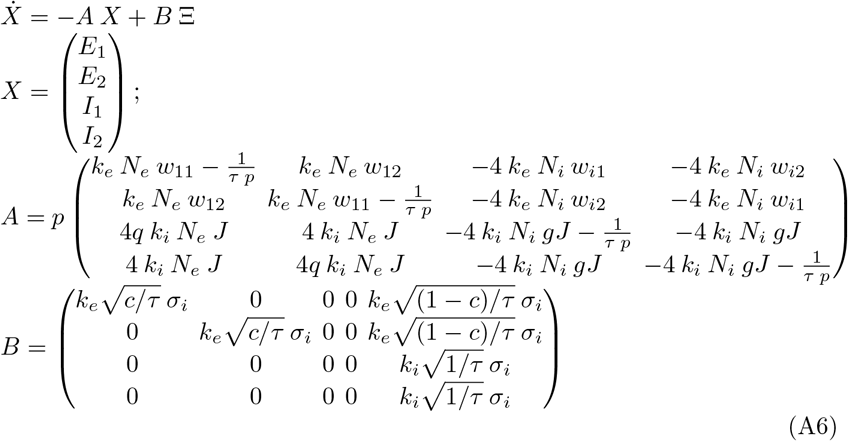

In this formalisation, the inhibitory weights from *I*_1_ and *I*_2_ to *E*_1_ and *E*_2_ are also dynamic and operate on a much slower time scale than the dynamics of the firing rates on the left side of (A5). To obtain the governing equations for individual weight dynamics, we assume that these weights which also play a role in shaping the dynamics of neuronal activities in equation A6 are piece-wise constant (separation of time scales).

To obtain the slow dynamics of the inhibitory weights onto the excitatory neurons (*w*_*i*1_ and *w*_*i*2_), as suggested in [24], the cross correlation function between the pre (inhibitory) and post (excitatory) synaptic firing rates, as well as the firing rates of the excitatory and inhibitory neurons at each moment in time are required:

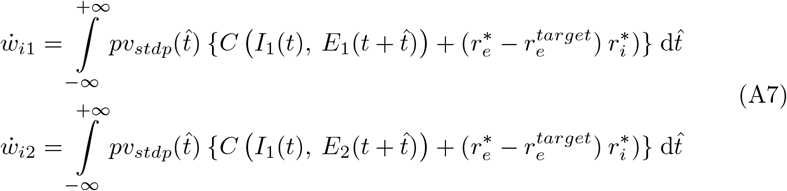

Where the symmetric stdp function which defines *pv_stdp_*(*t*) is exp(–|*t*|/*τ_stdp_*). The functions 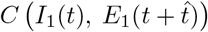 and 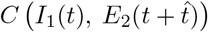 are the cross covariance functions between *I*_1_ and *E*_1_, and *I*_1_ and *E*_2_, respectively. The variables 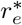 and 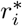 are the solutions for the average firing rates of the cells, which also depend on the weights *w*_*i*1_ and *w*_*i*2_, and are functions of time, but the homeostasis provided by the inhibitory plasticity ensures that the average firing rates of the excitatory cells in *E*_1_ and *E*_2_ converge the target value 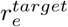. Due to this dependence on time, and also dependence of the rates and weights on one another, the whole set of equations (A7) and (A5) have to be solved self-consistently.

According to (A7), in order to understand the dynamics of weigh evolution for *w*_*i*1_ and *w*_*i*2_, we need to first evaluate the covariance functions between the excitatory and inhibitory average neuronal dynamics, and then multiply them by the exponential stdp function provided by *pv_stdp_*. It is known from [51] that for coupled Ornstein-Uhlenbeck (OU) processes, the expected covariance matrix can be obtained as follows

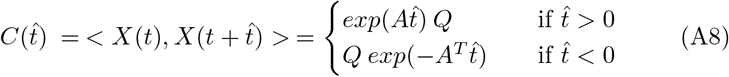

where *Q* is the solution to the following Lyapunov equation, which incorporates the correlation structure of the input in matrix B (defined in equation (A6)).

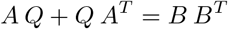

The challenge is to write the weight dynamics (A7) in a self-consistent way, such that it only depends on the weights of the network and the correlation structure in the input written as a function of coupling weigths. Since the right hand side of the equations in (A7) are integrals of exponential functions multiplied by a covariance function, we can evaluate the integral in the Laplace domain. For that, first we need to find the Laplace transforms of the covariance functions in equation (A8). Since the impulse response of the general system 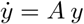 is *exp*(*At*) in time domain, and (*s***I** – *A*)^-1^ in the Laplace domain (*s* is the complex variable defined in the Laplace transform, and **I** is the identity matrix), the matrix exponentials in equation (A8) can easily be replaced by (*s***I** – *A*)^-1^ and (*s***I** + *A^T^*)^-1^, respectively.

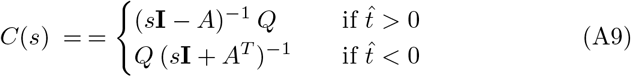

and the element on row *i* and column *j* of matrix *C*(*s*) will be represented by *C_i,j_*(*s*). The covariance function between the inhibitory populations and the excitatory populations for all positive and negative lags can therefore be achieved as following:

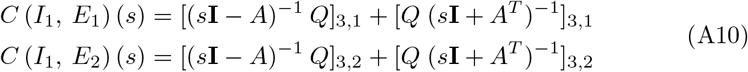

This can help to evaluate the integrals in equation (A7) because the integral of any function from *t* = 0 to *t* = ∞ can be calculated by evaluating that function at *s* = 0 in the (one-sided) Laplace domain. It is also known that the product of a linear system and an exponential function causes a shift in the Laplace domain. In other words, for the stdp function in the form of exp(– *t/τ_stdp_*), the integrals in equation (A7) in the Laplace domain should be evaluated at *s* = 1/*τ_stdp_*. Recruiting these mathematical tricks, equations (A8) and (A7) can be rewritten as

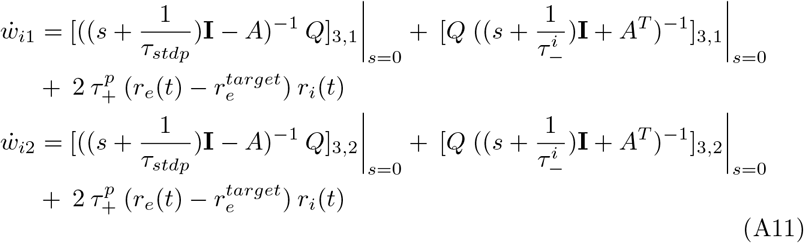

where *r_e_*(*t*) and *r_i_*(*t*) are evaluated in a recursive way and the whole system of equations above is solved in discrete domain. Starting from identical initial conditions for *w*_*i*1_(0) and *w*_*i*2_(0), and the corresponding solutions for *r_e_*(0) and *r_i_*(0), the weight dynamics of *w*_*i*1_(*t*) and **w*_*i*2_*(*t*) can be obtained iteratively. Since the matrices *A* and *A^T^* include the dynamical variables *w*_*i*1_ and *w*_*i*2_, we will end up with a set of coupled ODEs that describe the evolution of these variables. This can result in a state space plot for the dynamics of these weights.

### Table of parameters used in network simulations

Table of parameters for different network studies in the paper is as follows.

**Table A1.** Parameters unsed in network simulations for figures 1 to 7

| Variable | Explanation | figure 1 and 7 | figure 2 and 3 | figure 4 | figure 5 and 6 |
| --- | --- | --- | --- | --- | --- |
| $N_e$ | Number of E cells in each assembly | 800 | 3200 | 800 | 800 |
| $N_i$ | Number of PV neurons | overall 400 | overall 1600 | 200 per group | 200 per group |
| $J$ | EPSP amplitude for fixed connections (mV) | 0.04 | 0.02 | 0.04 | 0.04 |
| $w$ | scalar for within assembly weights | 2.5 | 2.5 | 2.5 | 2.5 |
| $g$ | Scalar for the fixed IPSP amplitudes | 10.0 | 10.0 | 10.0 | 10.0 |
| $\theta$ | Threshold for the membrane potential (mV) | 20.0 | 20.0 | 20.0 | 20.0 |
| $v_{reset}$ | membrane potential reset level (mV) | 10.0 | 10.0 | 10.0 | 10.0 |
| $v_{rest}$ | resting potential (mV) | 0.0 | 0.0 | 0.0 | 0.0 |
| $\tau$ | membrane potential time constant (ms) | 20.0 | 20.0 | 20.0 | 20.0 |
| $C$ | Capacitance of the membrane potential ($\mu$ F) | 250.0 | 250.0 | 250.0 | 250.0 |
| $p$ | probability of connection between E cells | % 10 | % 10 | % 10 | % 10 |
| $p_{ie}$ | probability of connections from E to PV | % 40 | % 40 | % 40 | % 90 |
| $p_{ei}$ | probability of connections from PV to E | % 40 | % 40 | % 40 | % 40 |
| $p_{ii}$ | probability of connections between PV cells | % 40 | % 40 | % 40 | % 40 |
| $\tau_{stdp}$ | time constant of PV to E stdp (ms) | 20.0 | 20.0 | 20.0 | 20.0 |
| $\eta_e$ | learning rate for E to E connections | supp: 0.000015 | NA | NA | NA |
| $\eta_e$ | learning rate for E to E connections | supp: 0.000005 | NA | NA | NA |
| $\eta_p$ | learning rate for PV to E connections | supp: 0.00005 | 0.00005 | 0.00005 | 0.00005 |
| $c$ | percentage of shared input from the total input | % 30 | % 30 | % 30 | variable |
| $\sigma$ | std of the external Poisson process (Hz) | $\sqrt{37500}$ | $\sqrt{60000}$ | $\sqrt{30000}$ | E cells: $\sqrt{37500}$<br>PV cells: $\sqrt{24375}$ |

## Supplemental Figures

### Network with log-normal E to PV distribution and excitatory Hebbian plasticity

**Fig. A1.**
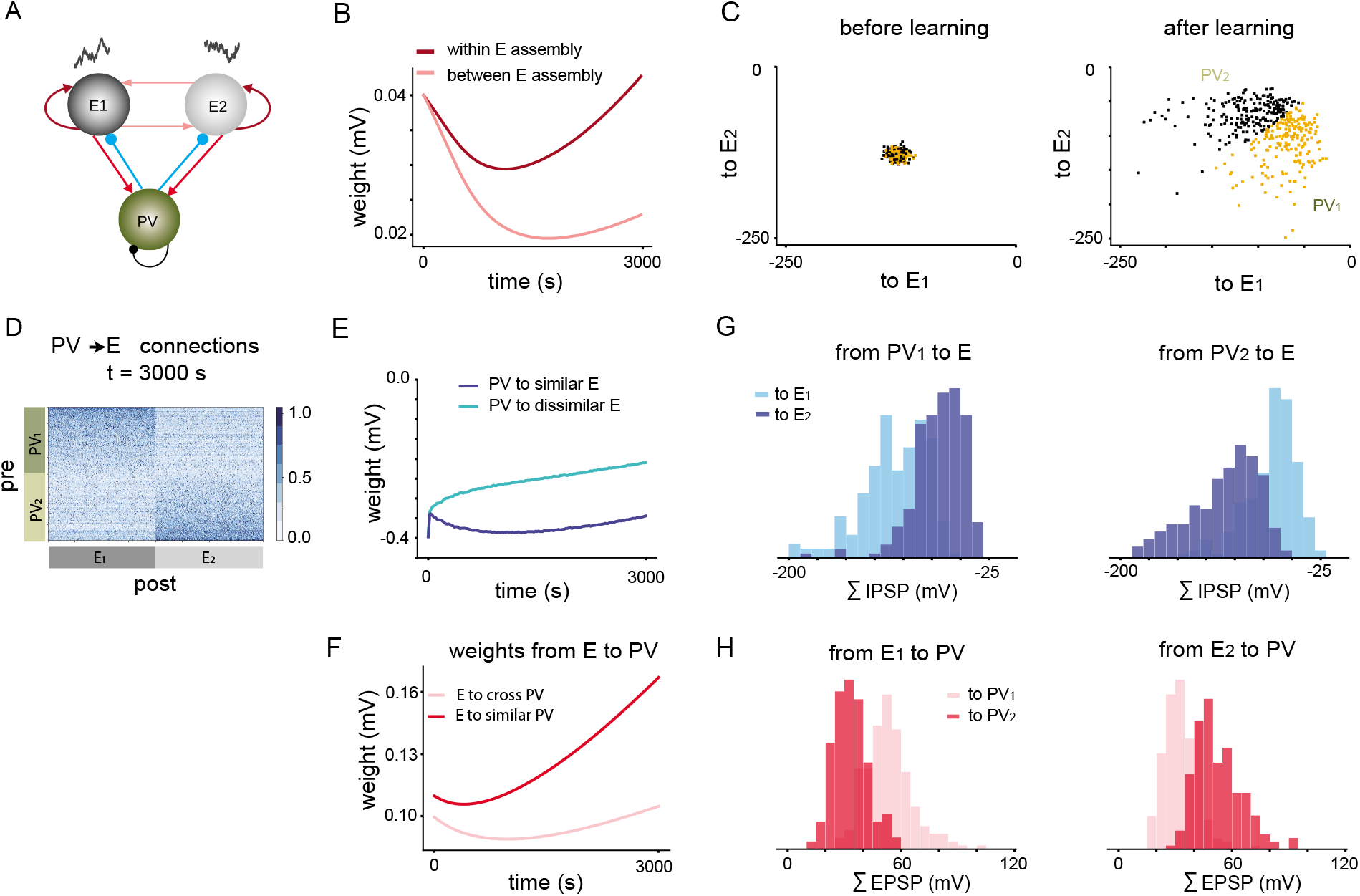
Emergence of tuned PV to E populations in a fully plastic network with fixed PV to PV weights. **A**: Network initially composed of two E assemblies with distinct shared input sources, and one PV population. **B**: Excitatory weight within and between E assemblies initially depress, which merely reflects the transient dynamics in the weight evolution. Weights within E assemblies are stronger than those between assemblies. **C**: Sum of IPSP projections of all PV cells to *E*_1_ and *E*_2_ assemblies (outgoing PV measure) define 2D vectors that are all very close to each another (left). After learning, some PV cells develop bigger total IPSP sums to one of the E assemblies (right). PV cells with bigger IPSP sum to *E*_1_ are labeled as PV_1_, and the rest of the PV cells are labeled as *PV*_2_ cells. **D**: Connectivity matrix for the connections from PV to E cells. **E**: Average weight evolution for the connections from PV_1_ to *E*_1_ (more negative weights indicate stronger connections) and to *E*_2_. **F**: Excitatory weights from E to PV cells follow a Hebbian STDP, and after turning on the plasticity mechanism, initially depress. However, *E*_1_ to PV_1_ weights become stronger as a function of time then *E*_1_ to *PV*_2_ weights. This, consequently, plays a role in assigning PV clusters. **G**: Distribution of PV to *E*_1_ and *E*_2_ assemblies indicate that PV_1_ cells projected more strongly to *E*_1_ cells, by virtue of their labeling. **H**: left: Strong projections from *E*_1_ to PV_1_ and relative weaker projections from *E*_1_ to *PV*_2_ (also by symmetry, right: strong projections from *E*_2_ to *PV*_2_ and relative weaker projections from *E*_2_ to PV_1_) are the main drive of PV tuning.

### Covariance structure for wide, narrow and delta networks

**Fig. A2.**
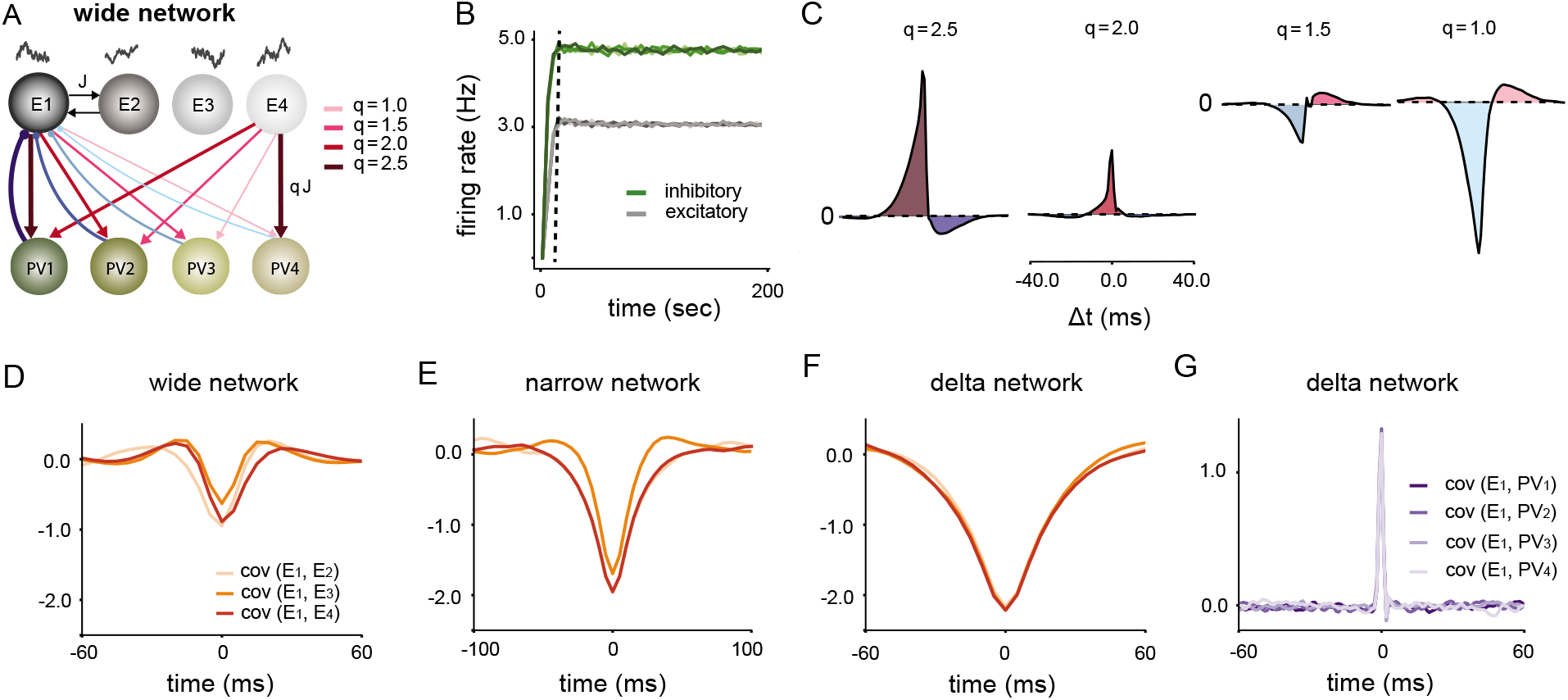
Tuned PV to E assemblies decreases competition between assemblies. **A**: Wide network and its corresponding results from B to D. **B**: Firing rates of the excitatory and inhibitory assemblies converge to their stationary rate very quickly. After a few ten seconds of simulation, the firing rates of all neurons reach a steady state value. Dashed line indicates the onset of stationary mean firing rate for the network. **C**: Product of the covariance functions in Fig. 4C with the symmetric STDP function results in a bigger net potentiation for *q* = 2.5. With decreasing values of q the integral becomes more negative which indicates depression of the inhibitory weight. *q* = 1.0 results in the most negative value (more depression). **D**: Covariance function between the excitatory assemblies are negative with a relatively smaller amplitude around the zero-lag value. **E**: For the narrow network, the negative amplitude for the cross-covariance function between the excitatory assemblies takes intermediate values. **F**: The largest negative amplitude for the cross-covariance functions belong to the network with delta distributed projections from E to PV cells. **G**: For the delta network, cross-covariance functions between *PV*_1_ and all other excitatory assembly firing rates are mainly positive and identical between assemblies.

### Effect of wide vs narrow E to PV weights in shaping the relationship between EPSP and IPSP

**Fig. A3.**
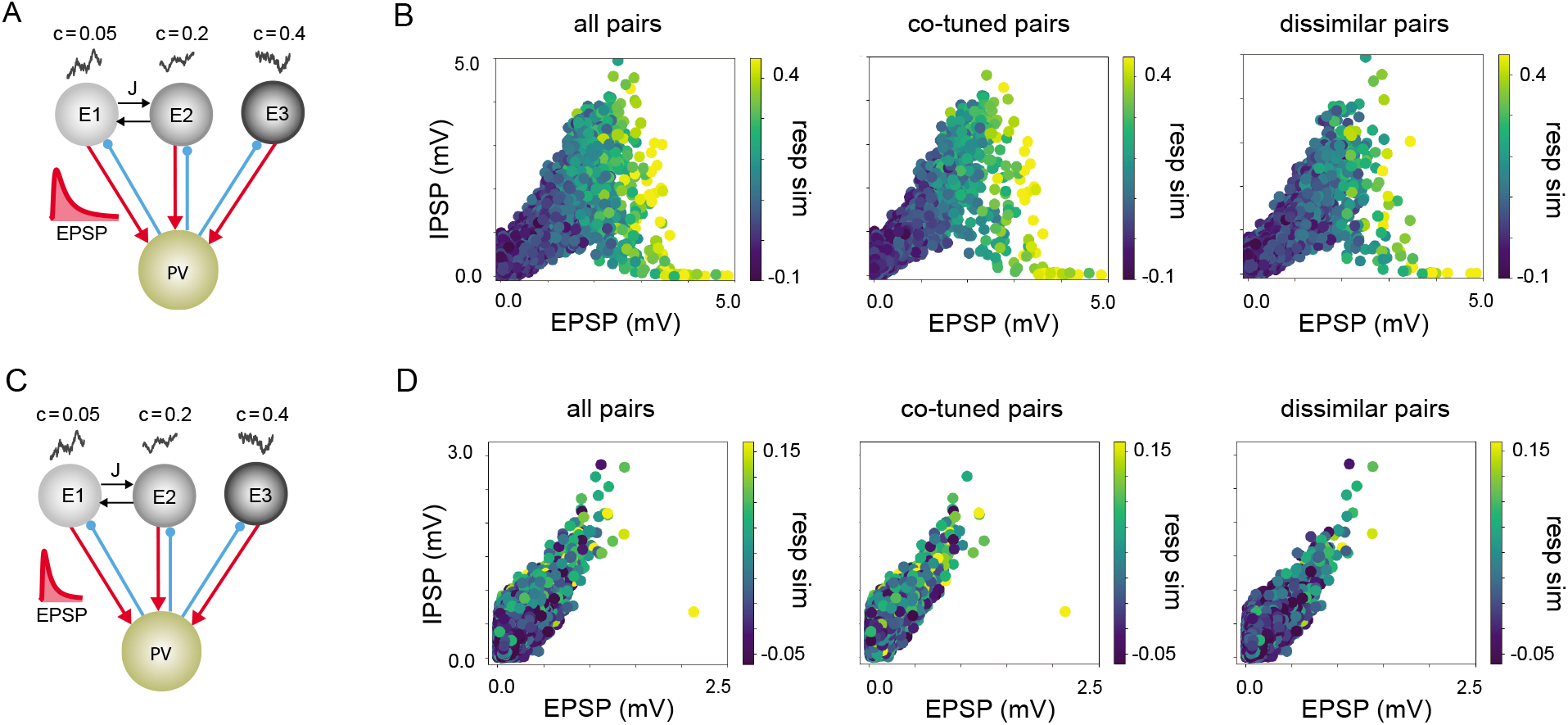
Effect of high and low variance in E to PV connection strength on the reciprocal PV to E connection weights. **A**: Network schematic for connections with high variance for the log-normal distribution. **B**: Scatter plot of EPSP and IPSP with pairwise response similarity as color codes for the network in A. Co-tuned and dissimilarly tuned pairs have been plotted separately. Very high values of EPSP result in decresed IPSP amplitudes for large values of EPSP. **C**: Network schematic for connections with low variance for the log-normal distribution. **D**: EPSP and IPSP relationship and response similarity between pairs of reciprocally connected PV and E cells for low variance connections follows a linear trend.

### Labeling PV cells based on their maximum summed IPSP projections onto E assemblies (outgoing PV measure)

**Fig. A4.**
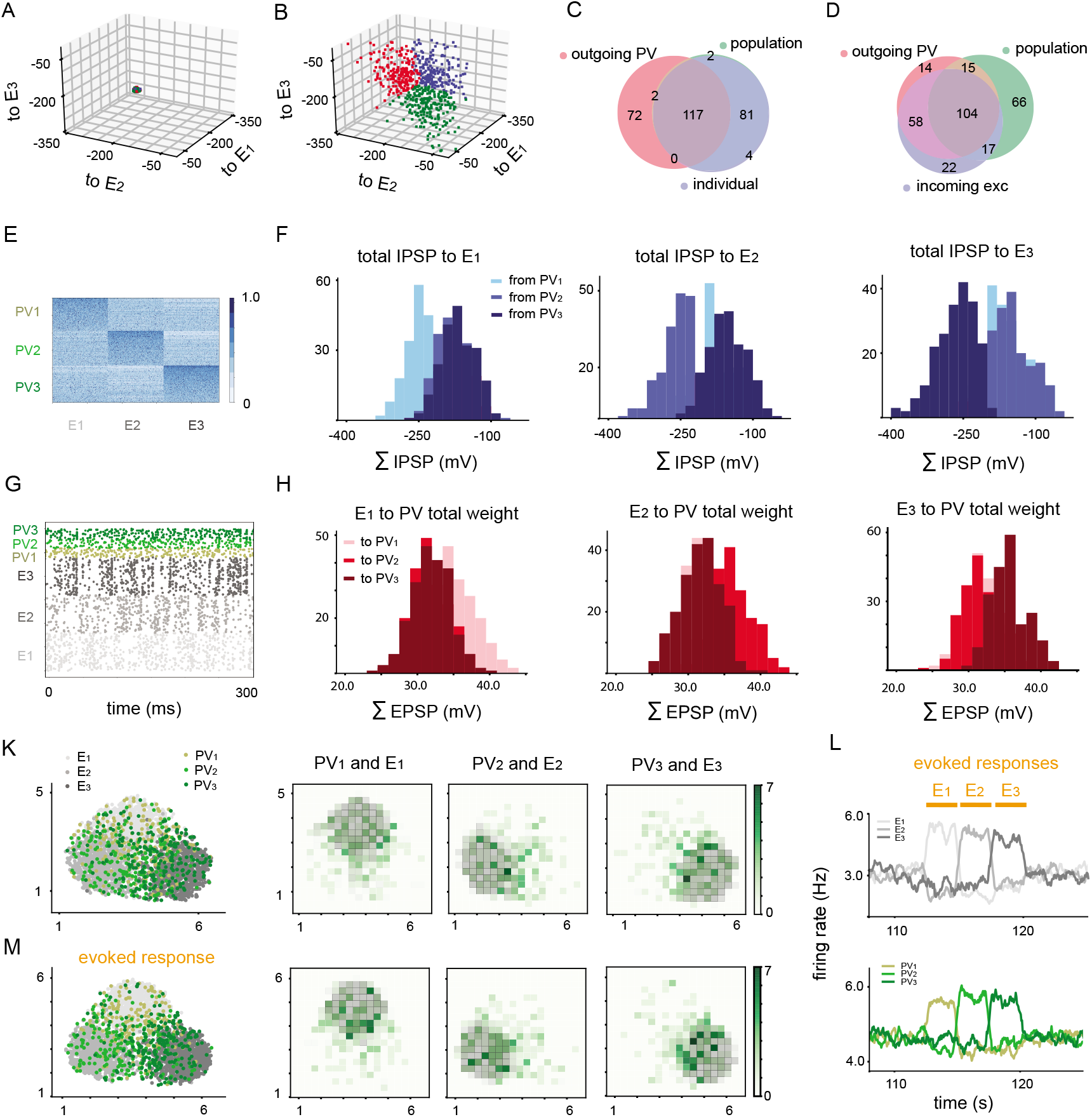
Labeling PV cells based on their maximum outgoing summed IPSP projections (outgoing PV measure) in the evoked state. **A**: Sum of the IPSP amplitudes of the projection weights from a single PV cell to all neurons in *E*_1_, *E*_2_, and *E*_3_ form one small 3-dimensional cluster of vectors, before plasticity operates. Colors are inherited from B after assignment of feature preferences. **B**: After plasticity is turned off, PV cells divide into 3 distinct groups with more preference to connect strongly to one of the excitatory assemblies: blue: preference to strongly connect to *E*_1_; red: preference to strongly connect to *E*_2_; green: preference to strongly connect to *E*_3_. **C**: Venn diagram for the labeling of PV cells that are assigned to PV_1_ based on 3 measures: outgoing PV, population similarity, and individual similarity. The overlap between the last two measures is almost %100. **D**: Venn diagram for the labeling of PV cells that are assigned to PV_1_ based on 3 measures: outgoing PV, population similarity, and incoming E. There is a large overlap between any two measures. **E**: Connectivity matrix from PV to E cells after labeling PV cells based on the outgoing PV measure. **F**: Distribution of the total IPSP weights from neurons in individual PV populations to *E*_1_ indicate that PV_1_ has the highest preference of connection to *E*_1_, hence these group of PV cells were labeled as PV_1_ (left). Similarly, PV_2_ (PV_3_) neurons have the strongest projections onto *E*_2_ (E_3_). **G**: Raster plot of neuronal activities for the last 300 ms of ongoing plasticity. **H**: Total EPSP projections from individual cells in *E*_1_ onto the assigned PV populations indicate that *E*_1_ had a stronger total projection weight onto PV_1_ neurons (left). A similar relation holds for other excitatory assembly projections: e.g summed EPSP weights from *E*_3_ onto PV_3_ were more skewed towards bigger values (right). **K,M**: 2-dimensional UMAP projection of the neuronal activities for 7.5 seconds of simulation time according to their distance in the high dimensional space for the spontaneous (K) and evoked (M) state. Planar density of co-tuned excitatory and PV cells are plotted separately for each feature defined by shared correlated input. **L**: Evoked responses of the E and PV subnetworks as a result of sensory stimulation of individual E assemblies.

### pairwise versus population measures can result in different conclusions for the tuning of PV cells

**Fig. A5.**
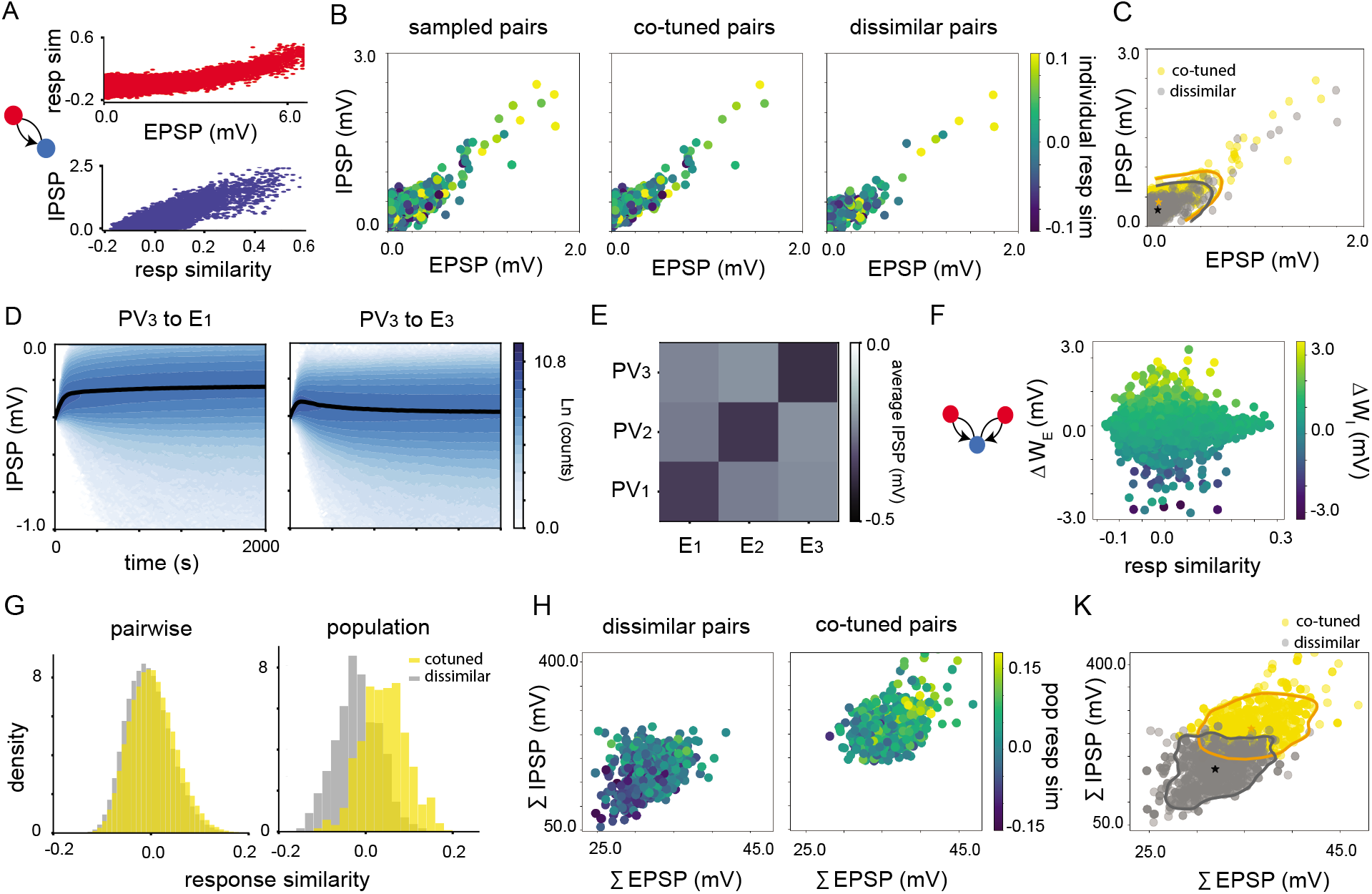
Pairwise and population measures for defining PV subnetworks. **A**: For reciprocally connected E and PV cells in the spontaneous state, response similarity increases with increasing values of EPSP. Strength of IPSP increases monotonically with pairwise correlation coefficient. **B**: left: EPSP and IPSP values for 2000 sample of reciprocally connected pairs. middle, right: Reciprocal EPSP and IPSP values for pairs belonging to similarly tuned populations and dissimilarly tuned populations. **C**: Distribution of cotuned and dissimilarly tuned pairs. Contour lines define area which contains %95 of the data for each group. Mean EPSP and mean IPSP for each group is indicated by a star (orange: cotuned group, gray: dissimilarly tuned group). **D**: Evolution of IPSP distributions from *PV*_3_ cells to *E*_1_ (oppositely tuned) and *E*_3_ (similarly tuned) cells as a function of time. **E**: Average connectivity matrix for the weights from PV subnetworks onto excitatory assemblies. **F**: Difference between the IPSPs for the triplets of E-PV-E as a function of cosine similarity between the excitatory cells and the difference between the projecting excitatory weights onto the shared PV cell. As Δ *W_E_* increases, Δ *W_I_* also increases. **G**: Density distribution of cosine similarity between pairs of E and PV cells belonging to similarly or dissimilarly tuned E assemblies are not distinguishable, however, if response similarity between individual PV cells and population responses of different E assemblies are considered, the distributions of the response similarities for the co-tuned and dissimilarly tuned PV cells become more separable. **H**: Summed IPSP of individual PV cell projection onto similarly (right) and dissimilarly (left) tuned E cells as a function of the sum of the EPSP received by the E assemblies. The colorbar defines cosine similarity of the PV cells with the average excitatory populations **K**: 2D scatter plots for cotuned an dissimilar pairs and contour lines defining regions in the space with %95 of the data for each case. Stars represent the mean values of the total EPSP and IPSP for each case. Summed IPSP as a function of summed EPSP on average takes bigger values when the sum of the EPSPs are large (neurons are similarly tuned).

### Covariance between population firing rates for the network with two assemblies

**Fig. A6.**
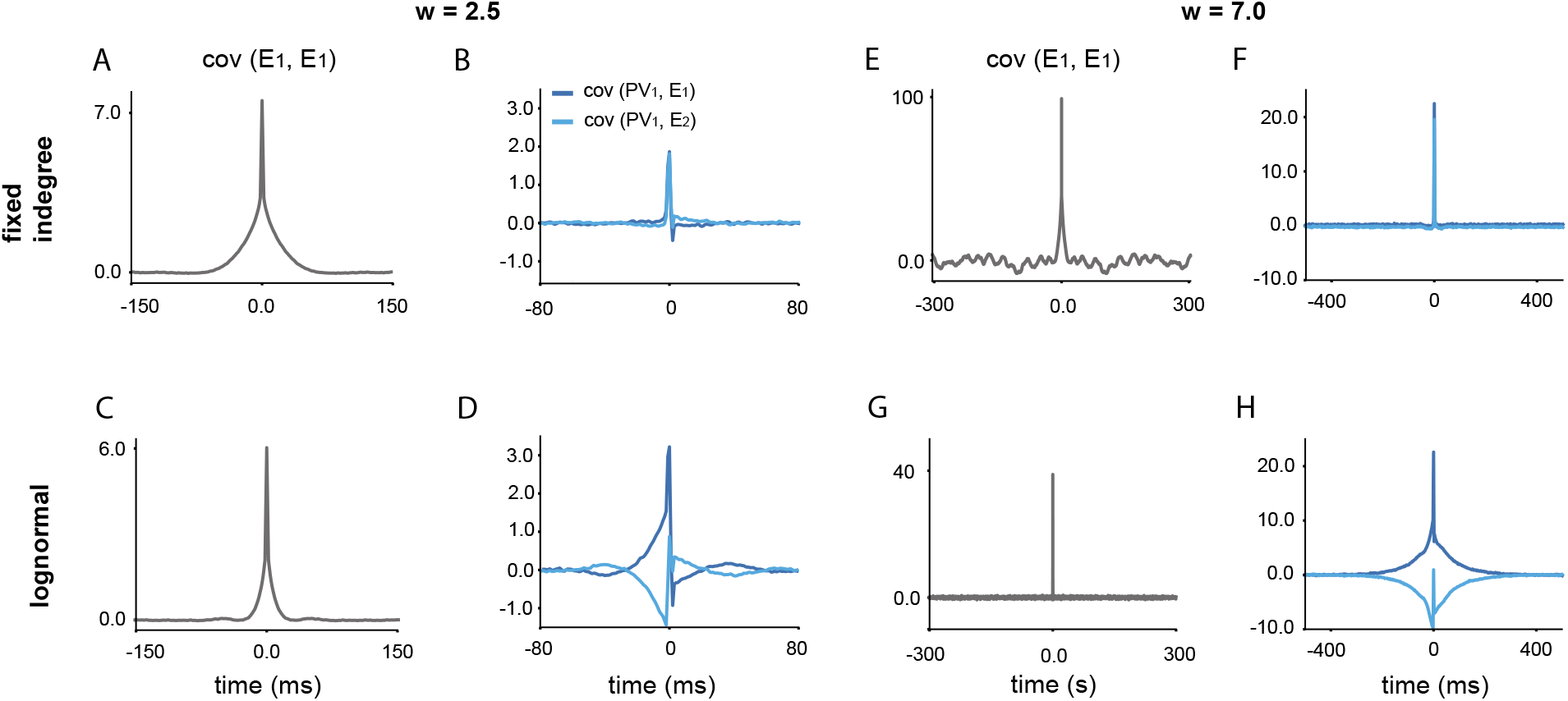
Covariance between subnetwork activities in Fig. 7. **A,C**: Auto-covariance function for the network with fixed in-degree and log-normal distribution for *w* = 2.5. **B**: For the network with fixed in-degree and *w* = 2.5, both covariance functions between one of the PV population activities, and the firing rate of *E_1_* and *E*_1_ are almost identical. **B**: For the network with log-normal E to PV distributions, *PV*_1_ and *E*_1_ are positively correlated, while *PV*_1_ and *E*_2_ are negatively correlated. **C**: For the fixed in-degree network at *w* = 7.0, the covariance functions between PV_1_ and *E*_1_ and *E*_2_ are strongly positive and identical. **D**: At *w* = 7.0, for the network with log-normal distribution, the covariance function between PV_1_ and *E*_1_ is strictly positive, while the covariance between PV_1_ and *E*_2_ is strictly negative. **F,H**: Auto-covariance function for the network with fixed in-degree and log-normal distribution for *w* = 7.0.

